# ABI1 regulates transcriptional activity of Androgen Receptor by novel DNA and AR binding mechanism

**DOI:** 10.1101/2023.05.26.542350

**Authors:** Baylee A. Porter, Xiang Li, Neeru Arya, Fan Zhang, Sonia H. Y. Kung, Ladan Fazli, Htoo Zarni Oo, Yinan Li, Kenneth Marincin, Konsta Kukkonen, Henna Urhonen, Maria A. Ortiz, Allysa P. Kemraj, Eva Corey, Xuesen Dong, Vladimir A. Kuznetsov, Matti Nykter, Martin E. Gleave, Gennady Bratslavsky, Alfonso Urbanucci, Dominique Frueh, Alaji Bah, Leszek Kotula

## Abstract

Transcription regulates key functions of living organisms in normal and disease states, including cell growth and development, embryonic and adult tissue organization, and tumor progression. Here we identify a novel mechanism of transcriptional regulation by an actin regulatory and signaling protein, Abelson Interactor 1 (ABI1). Using prostate cancer models, we uncover a reciprocal regulation between ABI1 and the Androgen Receptor (AR). ABI1 is a direct, androgen-regulated target; in turn, ABI1 interacts with AR and its splice variant ARv7, and co-regulates a subset of specific transcriptional targets. ABI1 directs transcription through transient yet well-defined interaction of its intrinsically disordered region with DNA. Clinical evaluation shows that the ABI1-DNA binding (through Exon 4 splicing) and ABI1-AR interaction are regulated during androgen deprivation therapy and prostate cancer progression, thus controlling tumor plasticity through connecting actin cytoskeleton and cellular signaling to transcriptional regulation. We propose ABI1 as epigenetic regulator of transcriptional homeostasis in AR-driven cancers.

**Statement of importance:** This study describes fundamental discovery in prostate cancer identifying novel mechanism of transcription by unique DNA binding mechanism involving actin cytoskeleton regulatory protein ABI1. ABI1-DNA binding activity predicts survival of prostate cancer patients. Moreover, we discover ABI1-AR reciprocal regulation that has far reaching implications for tumor plasticity and androgen-sensitive pathogenesis.

## Introduction

Epigenetics regulates cellular and organismal fate, overlaying the role of genetics in human disease and tumor evolution. This is particularly evident in Androgen Receptor (AR) driven cancers, in which targeting transcriptional regulation with androgen pathway inhibitors (ARPI) is a mainstay therapy. These treatments lead to the induction of transcriptional reprogramming, plasticity, and re-adaptation, which becomes a major challenge in effectively targeting therapy-resistant cancers such as castrate-resistant prostate cancer (CRPC) [1].

AR pathway inhibition leads to transcriptionally-induced plasticity. The insult from the ARPI therapy leads to clonal selection of tumor cell population with either reversible or irreversible states, resulting in heterogeneity of prostate tumors. A critical mechanism of prostate cancer (PCa) heterogeneity is the deregulation of transcriptional co-activators of AR that contribute to DNA binding specificity [2–4]. Here, we identify a novel regulatory mechanism of AR-mediated transcription by a cytoskeleton- and signaling-regulatory protein, Abelson Interactor 1 (ABI1).

ABI1 is a key component of the WAVE regulatory complex that promotes branched actin polymerization underlying cell motility and cell-cell adhesion [5]. The WAVE complex delivers signals from cellular receptors to downstream signaling proteins [6]. ABI1 regulates PI3 kinase as well as nonreceptor tyrosine kinases such as ABL and SRC [7, 8], sequestering the latter from activating STAT3 [5] and thus preventing non-canonical WNT pathway activation. The latter observation underscores ABI1’s tumor suppressor role as a signaling protein in advanced PCa, but no role for ABI1 in transcription or DNA binding has been identified.

Here, we define ABI1-AR reciprocal regulation involving two non-exclusive molecular mechanisms of ABI1-AR and ABI1-DNA interactions. These interactions are co-regulated during PCa evolution and clinical treatment promoting tumor plasticity and progression. We show that ABI1 binds AR via a multivalent interaction mechanism resulting in the co-liquid-liquid phase separation (LLPS) of the N-terminal intrinsically disordered regions (IDRs) of AR and ABI1 as well as the C-terminal SRC homology 3 (SH3) folded domain of ABI1. We also demonstrate that ABI1 binds chromatin through the homeobox homology region (HHR) located within the IDR of ABI1. The HHR binds dsDNA without inducing a disorder-to-order transition, thereby resulting in the formation of a “fuzzy complex” [9]. The interaction between HHR and DNA is modulated by the splicing states of HHR’s Exon 4, whose expression is associated with high-grade, advanced-stage PCa tumors and lower survival. Together, our results demonstrate that the ABI1-AR transcriptional program is altered during tumor progression and by anti-androgen treatments, suggesting that ABI1 is a key plasticity regulator in PCa by coupling its actin regulatory and signaling functions to transcriptional regulation. Our results provide new paths for biomarker and drug design discovery in PCa.

## RESULTS

### ABI1 is an androgen-regulated AR target modulated by clinical hormonal therapies

Changes in ABI1 expression during the natural progression of PCa [5] prompted us to evaluate AR-dependence of the ABI1 gene in the clinical PCa samples undergoing androgen deprivation therapy (ADT) (**Fig. 1A-B**). Immunoexpression analysis indicated that ABI1 is overexpressed in CRPC compared to primary/untreated tumors. Moreover, ABI1 expression is suppressed by Neoadjuvant Hormone Therapy (NHT) and in tumors with neuroendocrine PCa (NEPC) phenotype (**Fig. 1A**). Analysis of mRNA expression in PCa tumors from the TCGA PanCancer Atlas and Firehose/PRAD datasets indicated a positive correlation of ABI1 and AR expression (**Fig. 1C-D**) [10, 11]. We further confirmed changes in ABI1 expression in PCa PDX models representing hormone-sensitive primary PCa, CRPC, and NEPC; and confirmed overexpression of ABI1 in CRPC and downregulation in NEPC models (**Fig. 1E-F**). Similarly, Abi1 expression was also downregulated in the prostatic epithelium of castrated mice (**Fig. 1G-H**). In summary, analysis of ABI1 levels in PCa tumor and mouse tissue indicates a positive correlation between ABI1 levels and AR activity.

**Figure 1.**
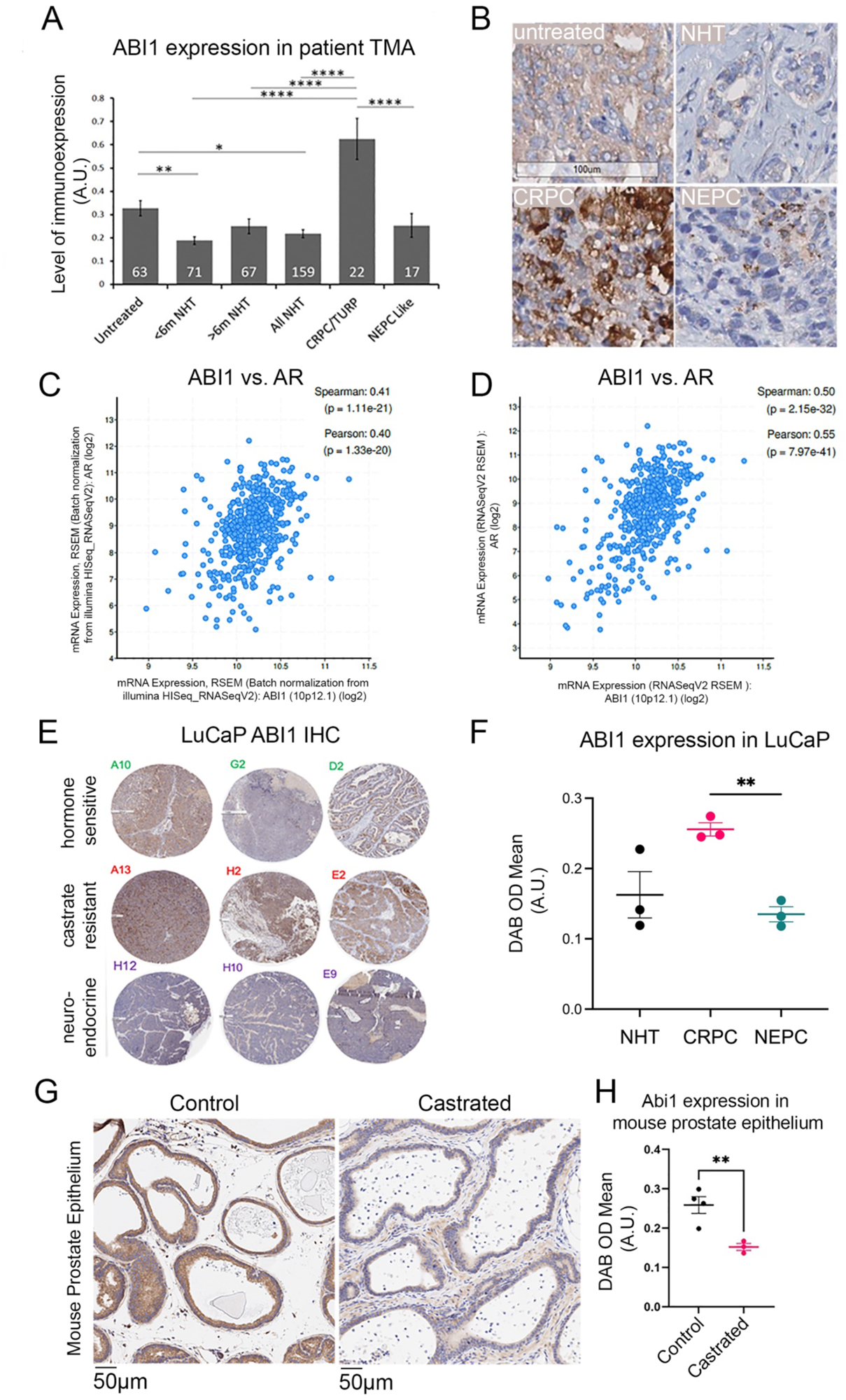
Correlation of ABI1 expression and the AR pathway activation levels during PCa tumor progression and hormonal treatment. **A**, Analysis of ABI1 expression (IHC with ABI1 antibody, quantified by digital imaging) in VPC TMA cohort of neoadjuvant hormone treatment (NHT) of PCa tumors. Numbers within bars show the total number of cases analyzed for each group, *, p<0.05; **, p<0.01; ****, p<0.0001. **B**, representative IHC images from TMA analysis in panel **A**. **C-D**, Correlation of ABI1 and AR mRNA expression in patient samples, in TCGA samples, PanCancer Atlas (n=494) **C**; or in Firehose/PRAD cohort, (n=500) **D**. **E**, ABI1 immunoexpression in TMA samples of LuCaP PDX cell lines, letters in left upper corners of images define position on TMA slides (HS, LuCaP 35/81/136; CRPC, LuCaP 35CR, 81CR, 136CR; NEPC, LuCaP, 77CR/86.2/93/145.1/145.2). **F** Quantification of ABI1 expression in **E**, n=3,** p<0.01. **G**, Abi1 immunoexpression in mouse prostate epithelium was decreased after 4 weeks of castration compared to non-castrated controls quantified in **H**, n=3, **p<.0098. *Abbreviations: HS, Hormone sensitive; CRPC, castrate resistant prostate cancer; TURP, transurethral resection of the prostate; NEPC, neuroendocrine prostate cancer; IB, immunoblot detection of indicated antibody target*.

It was important to determine whether *ABI1* is a *bona fide* AR-regulated gene as suggested by the expression data. The hormone-sensitive cell line LNCaP showed increased expression of ABI1 upon treatment with synthetic androgen R1881 while treatment with enzalutamide, an AR inhibitor, decreased ABI1 expression (**Fig. 2A-B**). Investigation of ChIPseq datasets of hormone-dependent cell lines, including LNCaP [12, 13] and VCAP [13–15], revealed the presence of an intronic AR-binding site (ARBS) within the *ABI1* gene (**Fig. 2C**) and the presence of active methylation marks (H3K4m1/3) and H3K27ac upstream, indicating active transcription and regulatory regions (**Supplementary Figure S1)**. ChIP-qPCR of the ARBS within the second intron of the *ABI1* gene increased AR binding thus confirming regulation of ABI1 expression (**Fig. 2D**). In the same LNCaP cells overexpressing AR [16] we showed that increased AR binding at the ARBS within the *ABI1* gene correlated with the increased *ABI1* gene mRNA expression (**Fig. 2E**). These data indicate direct transcriptional regulation of ABI1 by AR.

**Figure 2.**
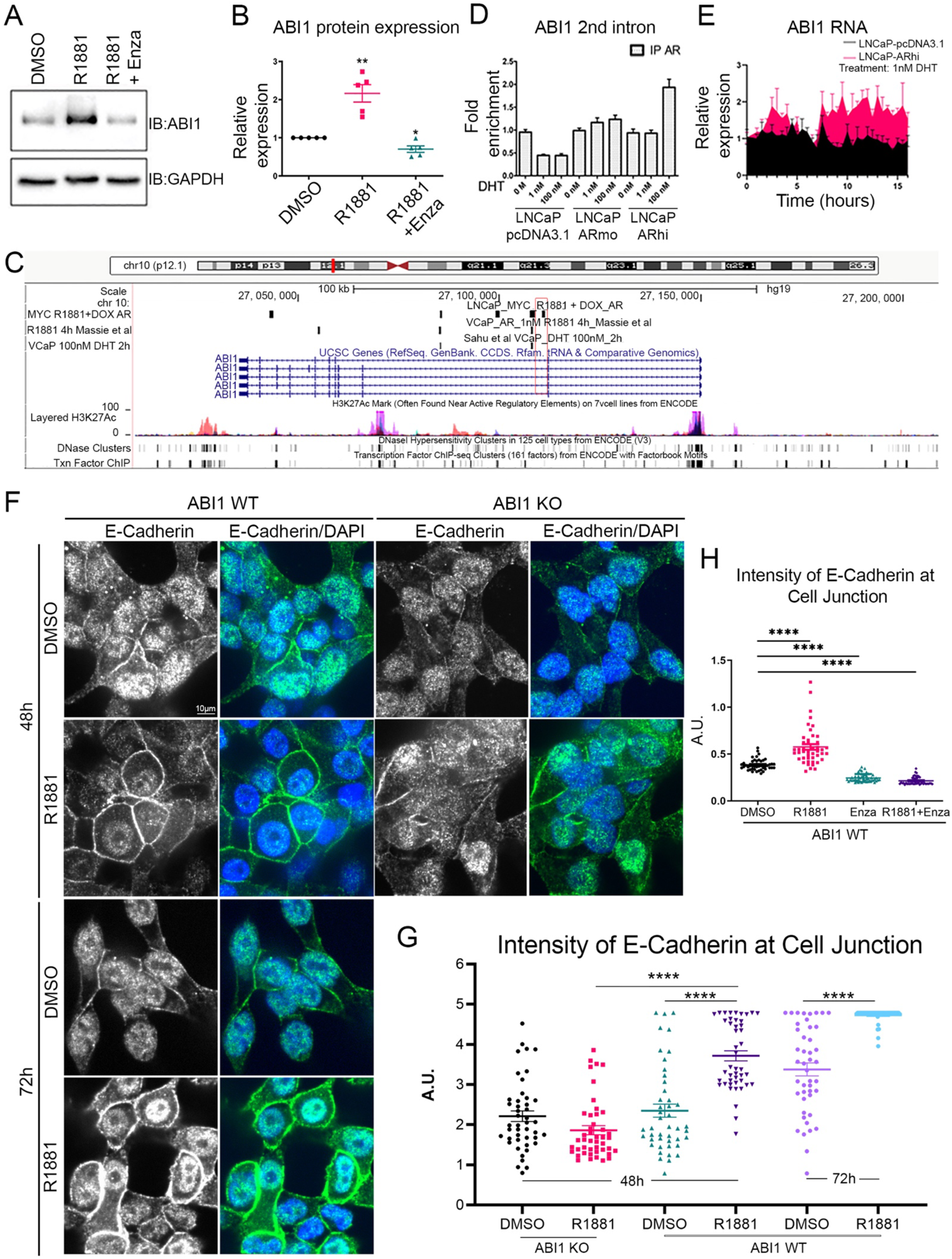
Transcriptional regulation of ABI1 and promote tissue integrity through ABI1-E-Cadherin axis. **A.** Expression of ABI1 is enhanced with synthetic androgen, R1881 (1 nM), but downregulated with AR inhibitor Enzalutamide (Enza) (5 μM). **B**. Quantification of ***A***, DMSO control versus R1881 (**p<0.007) and control versus R1881 + Enza (*p<0.02); n=3. **C.** UCSC genome browser-based diagram depicting AR binding site within *ABI1* gene. Black lines indicate binding events at specific locations across chromatin. Red box indicates AR binding locations on *ABI1*. **D**, ChIP-qPCR assays indicates a fold enrichment of AR binding to the 2nd intron binding site within *ABI1* upon overexpression of AR and DHT treatment in the LNCaP cells. **E**, Induction of ABI1 mRNA expression upon AR overexpression, LNCaP-AR vs. LNCaP-empty vector in DHT (1 nM) treated cells. **F**, Enhanced E-cadherin localization to cell-cell junction as functional readout of ABI1 at 48h and further at 72h (lower panels) compared to control but disrupted in ABI1 KO cell lines (right panels), quantified in ***G***. **H**. Inhibition of E-cadherin immuno-intensity at cell-cell junctions following Enza treatment alone or following R1881, vs. control (DMSO) (see **Supplementary Figure S1B** for representative images). Line intensities were quantified from 50 junctions per treatment; n=3; p<0.005.

We then investigated whether AR stimulation impacts a known function of ABI1 in promoting cell-cell adhesion. Previous studies have shown that ABI1 regulates cell-cell adhesion through E-cadherin localization at cell junctions through downstream WAVE complex formation [17]. AR stimulation with R1881 in LNCaP cells promoted E-cadherin levels at cell junctions in a hormone sensitive manner (**Fig. 2F, H-I**). E-cadherin accumulated at 48-, and 72-hours post-hormone stimulation in presence but not in absence of ABI1, indicating that AR-dependent regulation of E-cadherin at cell junctions is mediated by ABI1 (**Fig. 2G, J).** Collectively, these data indicated that ABI1 regulation of cell-cell adhesion is AR regulated and androgen-dependent.

### ABI1 interacts with the full-length AR and its splice variant ARv7 in prostate cancer cells and tissues

Since both AR [18] and ABI1 [19] can reside in the cytoplasm and the nucleus, we investigated whether these proteins could interact in these subcellular compartments. Protein sequence analysis of AR and ABI1 indicated a potential binding region between the proline rich motifs within the N-terminal Intrinsically Disordered Region (N-IDR) of AR and the C-terminal SRC homology 3 (SH3) domain of ABI1 (**Fig. 3A**). Using co-immunoprecipitation, we demonstrated that ABI1-AR interacted across multiple PCa lines (**Fig. 3B-F**). In 22Rv1 cell lines we observed the interaction of ABI1 not only with the full-length AR, but also with the truncated AR splice variant ARv7 (**Fig. 3E**), notoriously associated with emergence of CRPC [20]. By expressing either the full-length AR or ARv7 in the AR-negative cell line PC3 we further confirmed that ABI1 co-immunoprecipitated with both AR forms independently (**Fig. 3F**) which is consistent with the conservation of the PXXP-rich region in the ARv7 mutant (**Fig. 3A**). To examine whether the ABI1-AR interaction is SH3 domain-dependent, we performed experiments re-expressing ABI1 in the genetically modified LNCaP cells where the endogenous *ABI1* expression was disrupted by CRISPR and replaced with recombinant ABI1. This allowed us to probe ABI1 variants with mutations in SH3 or lacking SH3; here, we used the specific isoform 2 of ABI1 (ABI1 Iso2), which is one of the dominant Abi1 isoforms expressed in prostate tissue [21]. Co-immunoprecipitation experiments in recombinant LNCaP lines expressing ABI1 Iso 2 variants showed that the interaction between AR and ABI1 was partially abolished for an ABI1 Iso2-W485N mutant, lacking a critical residue within the SH3 domain [22]), or completely abolished in the mutant lacking the SH3 domain (ABI1-Iso2-ΔSH3) (**Fig. 3G-H**). These observations indicated that the interaction between AR and ABI1 is dependent on the SH3 domain.

**Figure 3.**
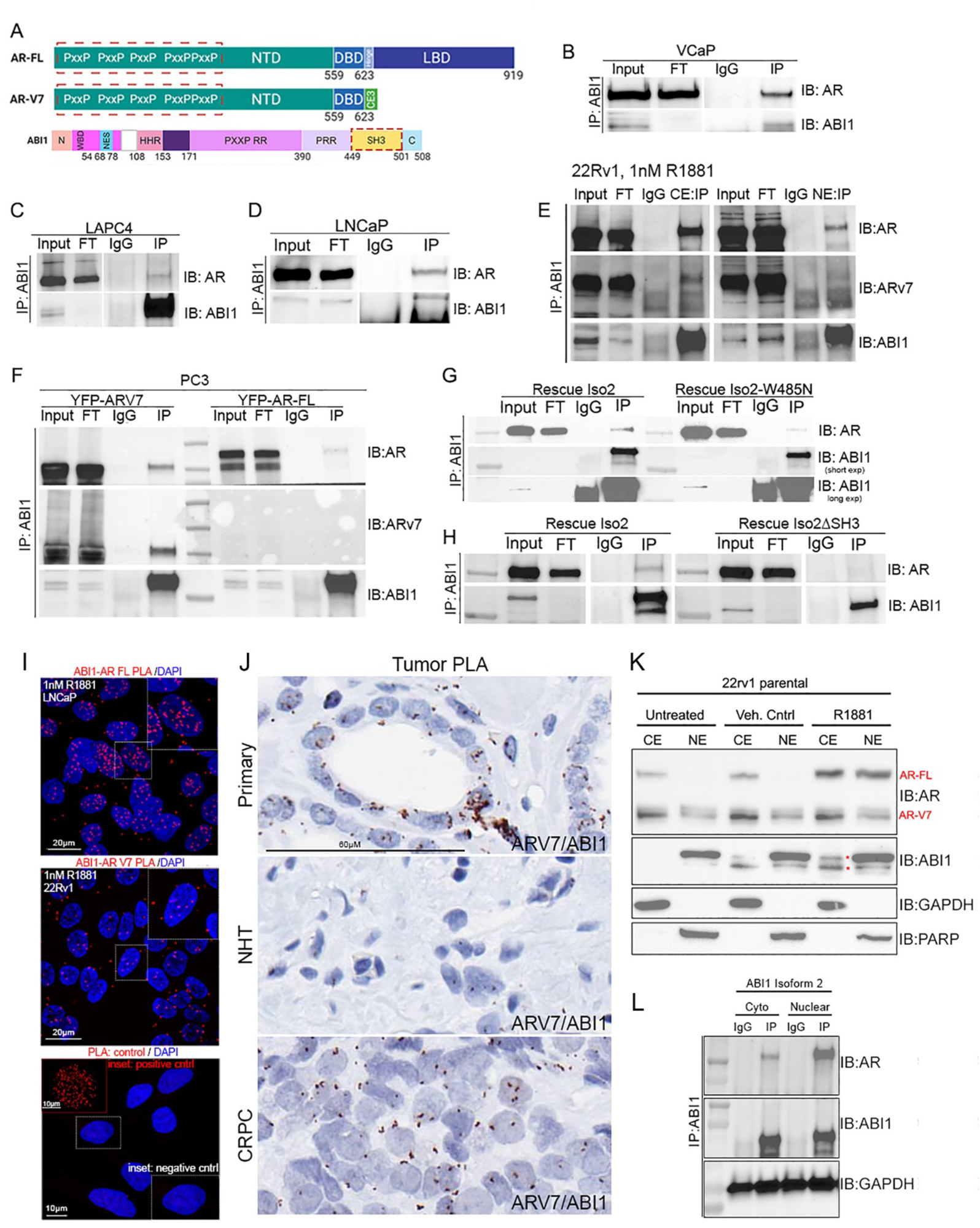
ABI1-SH3 domain interacts with AR-FL/ARV7 *in vitro* and PCa cells and tumors. **A**, Protein domain map of AR-FL/V7 and ABI1 with red boxes indicating their respective binding domains. **B**, Co-IP of ABI1 and AR in naïve VCaP, LAPC4 (**C**), LNCaP (**D**), 22Rv1 (**E**) cell lines indicating the interaction of ABI1 with the full-length AR (AR-FL) (all cell lines), and a truncated version (AR-V7) in 22Rv1 following the treatment with 1 nM R1881. **F,** Co-IP of ABI1 and AR-FL/ARV7 in a PC3 ABI1 isoform 2 (Iso2-WT) cell line transfected with AR-FL or ARV7. **G-H**, Co-IP of ABI1 and AR in LNCaP CRISPR ABI1 KO cells transfected with ABI1-WT, (Iso2-WT), or SH3 domain binding mutant (Iso2-W485N) (**G**), or SH3 domain deleted mutant (Iso2-ΔSH3) (**H**) showed decreased or no observable binding to AR vs. WT control, respectively. **I.** AR-FL/AR-V7 and ABI1 PLA interactions indicated by red color puncta, counterstained with DAPI in LNCaP cell line (*top panel*); 22Rv1 cell line (*middle panel*); positive control (*bottom panel inset*); negative control (*bottom panel*). Cells were incubated in charcoal stripped serum for 5 hours followed by 1 nM R1881 treatment overnight. **J.** AR-V7 and ABI1 PLA interactions (brown puncta) in primary, neoadjuvant hormone therapy treated, and castrate-resistant prostate cancer patient tumors. **K**, Immunoblot detection of AR and ABI1 protein expression in subcellular fractions of 22Rv1 cells treated with vehicle control or 1 nM R1881 for 24 hours. Loading controls include GAPDH (cytoplasmic fraction) and PARP (nuclear fraction). **L**, Co-immunoprecipitation of ABI1 and AR in subcellular fractions of LNCaP cells transfected with the recombinant ABI1 isoform 2. GAPDH, loading control. \**Abbreviations: NHT, neoadjuvant hormone therapy; CRPC, castrate resistant prostate cancer; CE, cytoplasmic fraction; NE, nuclear fraction; Cyto, cytoplasmic fraction; Nuclear, nuclear fraction; IgG, immunoglobulin G (negative control); IP, immunoprecipitation and antibody immunoprecipitated for; IB: antibody probe for on immunoblot; AR-FL: AR full length, AR-V7: AR splice variant 7. Red asterisks indicate ABI1 immunoreactive bands*.

### ABI1 interacts with AR through a phase separation mechanism

To understand the biochemical nature of ABI1-AR protein interactions we investigated a LLPS state [23] between these two proteins *in vitro*. AR was shown to form transcriptionally active biomolecular condensates, and while C-terminal folded DNA- and Ligand-binding domains of AR contribute to LLPS formation, the PXXP-containing N-IDR plays the predominant role [24, 25]. Using computational biology analyses, we found that ABI1 is predominantly disordered with only one structured SH3 domain at the C-terminus and an helical region shown to bind WAVE at the N-terminus [26] (**Supplementary Figure S2A-B** [27]). Interestingly, the N-IDR of ABI1 also contains multiple PXXP motifs, suggesting that both proteins may compete for the same ABI1-SH3 domain. Using recombinant proteins, we established conditions under which the N-terminus of AR can phase separate individually as demonstrated [28] or together with the full-length ABI1 (**Supplementary Figures, S3-4)**. This supports the hypothesis that multivalent interactions between AR and ABI1 can mediate the assembly of LLPS droplets *in vitro* and potentially form condensates in cells consistent with the proposed mechanisms [29].

To visualize the ABI1-AR interaction in prostate cancer cells we used proximity ligation assay (PLA) using AR, ARv7, and ABI1-specific antibodies. We observed the positive signal of ABI1-AR complexes in the nuclei of LNCaP cells treated with R1881 (**Fig. 3I**), and ABI1-ARv7 in 22Rv1 cells (**Fig, 3I**). Importantly, we also observed the positive PLA signal in clinical specimens of primary PCa, NHT, and CRPC tumors for ARV7-ABI1 complexes (**Fig. 3J**). These observations are consistent with ABI1 interacting with activated AR, or its constitutively active ARv7. To validate the ABI1-AR interaction using biochemical assays we performed subcellular fractionation followed by co-immunoprecipitation using LNCaP or 22Rv1 cells and confirmed the presence of ABI1-AR complexes in both nuclear and cytoplasmic compartments (**Fig. 3E, K, L**).

### ABI1 binds chromatin and is a transcriptional co-regulator of AR

Having established that AR and ABI1 complexes can phase separate and generate potentially transcriptionally active condensates in the nucleus of PCa (**Fig. 3I-J**), we investigated the nuclear functions of ABI1 in co-regulating AR-transcriptional activity. We first observed that the nuclear fraction of AR is decreased in ABI1-KO cells and rescued in cells expressing the wild-type ABI1 Iso2, but not in the ABI1 SH3 domain mutant, Iso2-W485N that cannot bind to AR (**Fig. 4 A-B**), suggesting that the presence of nuclear ABI1 is important for nuclear AR localization. We also observed that loss of ABI1 in LNCaP cells reduced several canonical AR-target gene expressions such as *KLK3*, *FKBP5* and AR, but did not affect *TMPRSS2,* another known AR target (**Fig. 4C**). Disruption of ABI1 also significantly reduced the expression of the *WASF1* gene encoding for a canonical WAVE complex component. Expressing ABI1-Iso2 recovered WASF1 expression but the two SH3 domain mutants did not (**Fig. 4C**). Importantly, *KLK3*, *FKBP5* and *AR* gene expression was enhanced in ABI1-Iso2 expressing cells. These results indicated that ABI1 contributes to AR transcriptional activity, and that its SH3 domain may be involved either through its interaction with AR, and/or its impact on AR nuclear localization.

**Figure 4.**
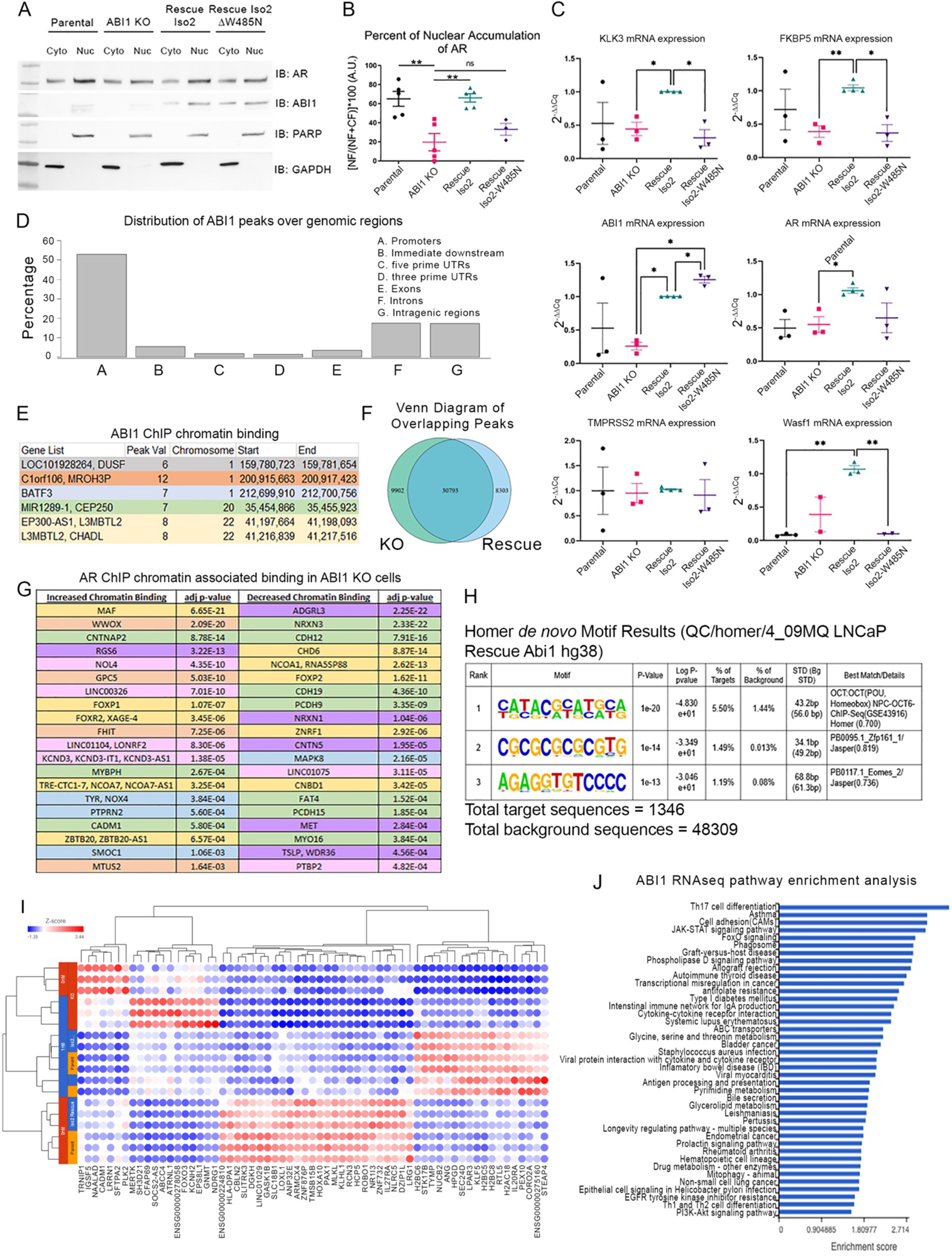
ABI1 regulates AR transcriptional program through specific chromatin binding. **A,** Immunoblot detection of AR, ABI1, PARP, and GAPDH in subcellular fractions of LNCaP cells: parental, ABI1 KO, ABI1 isoform 2 (Iso2), ABI1 isoform 2 SH3 domain mutant (Iso2-W485N). **B**, Quantification of AR in ***A***, indicates decreased nuclear localization of AR in ABI1 KO cells vs. cells rescued with ABI1 WT (Iso2) but not with the SH3 binding mutant W485N, (Iso2-W485N). Relative protein expression was quantified from densitometry measurements and normalized to fraction specific loading control either cytoplasmic fraction, GAPDH or nuclear fraction, PARP after subtraction of background signal. % Nuclear accumulation = [Nuclear Fraction/(Cytoplasmic fraction + Nuclear fraction)] * 100; n=3, **p<0.01. **C**, Transcriptional activity of known AR targets such as *KLK3, FKBP5,* and novel target gene *Wasf1*(Wave1) but not for *TMPRSS2* is reduced in cells lacking ABI1 (ABI1 KO) and in cells expressing SH3 binding mutant W485N, (Iso2-W485N). Scatter plots represent ΔΔCq mRNA expression collected using TaqMan Probe qPCR assay for known target genes of AR; n=3, *p<0.05, **p<0.01. **D.** Distribution of ABI1 peaks over genomic regions (as indicated) from ABI1 ChIP-seq data wherein ABI1 predominantly binds promoter regions suggesting ABI1 may play a role in transcription factor recruitment. **E**, Top AR-overlapping genes regulated by ABI1. **F**, VENN Diagram of “KO” represents overlapping peaks of AR ABI1 KO ChIP replicates 1 and 2 and “Rescue” represents overlapping peaks of AR ABI1 Rescue ChIP replicates 1 and 2. Unique peaks identified between samples were, KO=9902 and Rescue=8303 whereas common peaks= 30793. **G**, AR ChIP-seq data set illustrating 20 genes that have increased (left) or decreased (right) AR chromatin binding identified using genomic regions containing 1 or multiple overlapping Intervals and the associated FDR or “padj” cutoff for significance FDR < 0.001 based on DESeq2 analysis of 50kb. **H.** De Novo Homer Motif analysis from ABI1 ChIP seq samples indicated log ranked p-values in order of significance using Homer Motif analysis software. **I**, Hierarchal clustering map from RNAseq data for ABI1-Iso2-WT/KO cell lines (Iso2 rescue, or KO) shows 4 groups of genes positively and negatively regulated by ABI1, in the presence (0 nM) or absence of androgen treatment, R1881 (1nM). ABI1 negatively regulates gene expression of the first seven genes and treatment with AR canonical ligand further induces downregulation when ABI1 is present. **J**, Pathway enrichment analysis from RNAseq data shows that ABI1 is involved in pathways such as cell adhesion, JAK/STAT, FoxO, transcriptional misregulation in cancer. *Labels: green, cytoskeleton/adhesion; yellow, DNA binding; blue, cytosolic component; pink, RNA/splicing factor; orange, tumor suppressor; purple, signaling molecule*.

Given the effect of ABI1-Iso2 on the transcription of specific target AR genes we performed ChIP-seq to assess the ability of ABI1 to bind chromatin. ABI1 ChIP-seq in the ABI1-Iso2 LNCaP cells mapped binding predominantly to promoter regions (**Fig. 4D**). To determine the potential collaboration of ABI1-AR in transcriptional activity, we also examined whether the AR DNA binding profile could be modulated by ABI1. Data from the AR ChIP-seq profile of the same cells displayed AR chromatin binding sites in proximity of genes that had ABI1 bound at promoter sites (**Fig. 4E-F**). Most importantly AR ChIP-seq profiles in ABI1-KO and ABI1-Iso2 rescued LNCaP cells showed that the disruption of ABI1 resulted in reprogramming of ∼20% of AR chromatin binding. The loss of ABI1 resulted in chromatin occupancy alteration for AR in proximity of a subset of genes coding for cell adhesion proteins, chromatin remodeling complexes, Forkhead family of proteins, and other transcription factors (**Fig. 4G**). Motif sequence analysis using HOMER identified top ABI1 binding DNA sequences including the homeobox gene binding motif (**Fig. 4H**). These data demonstrated that ABI1 can bind specific DNA *in vivo* i.e., in cells.

RNA sequencing (RNA-seq) data from ABI1-KO and ABI1-Iso2 LNCaP cells indicated several pathways that were either positively or negatively regulated by ABI1 in response to androgen treatment (**Fig 4I**). These pathways include JAK/STAT, FOXO, and cell adhesion. Integration of ABI1 ChIP-seq data with RNA-seq data revealed correlations of chromatin binding with modulation of gene expression coding for multiple ABI1 associated complexes (CRK, CYFIP2, STAT3/5A), known EMT regulators (SMAD6, SMARCA4, SOX13, FZD1/8), microtubules (MAP9), transcriptional initiation complex/chromatin remodeling complexes (MED31, CHD4/9), and AR-ABI1 dually regulated genes (FKBP4, L3MBTL2, MED31, NCOR2, NCOA2/3/7, GLI3, PARP1/8) (**Fig 4J**). Overall, these results indicate that ABI1 binds to chromatin, and it is a transcriptional co-regulator of AR.

### ABI1 binds DNA through a binding mechanism involving the HHR region

ChIP-seq data indicated the potential for direct binding of ABI1 to DNA. Moreover, the analysis of ABI1-dependent expression profiles revealed the regulation of several homeobox transcriptional factors by ABI1 including HOXB4 [5] that has a weak homology to the ABI1 homeobox homology region or HHR (**Supplementary Figure 5**). Apparently, ABI1-HHR includes a 5 amino acid region encoded by alternatively spliced Exon 4. To determine the DNA-binding mechanism of ABI1 we designed several double stranded (ds)DNA probes derived from the CHIP and ABI1 expression analysis; representative dsDNA probes are listed in **Supplementary Material, Table 1**. We also used the ABI1-HHR lacking Exon 4 sequence (ABI1-HHR-DelEx4). An *in vitro* binding assay with dsDNA probes indicated binding affinities in the low micromolar range for dsDNA probes BATF3, EP300-AS1, MIR1289-1, C1orf106, L3MBTL2 and HOXB4. (**Fig. 5A-B**). The recombinant ABI1-HHR polypeptide spanning amino acid residues 93-170 of ABI1 retained similar DNA binding affinity as determined for the full-length ABI1. Assay results using DNA mutants demonstrated lower affinities for the 2-residue-mismatched dsDNA HOXB4 probe, and for the ABI1 HHR lacking Exon 4 (ABI1-HHR-DelEx4) (**Fig. 5C**). As the DNA mismatch introduces both sequence and shape changes into dsDNA, these data indicated the binding specificity of ABI1-HHR to HOXB4, as well as the critical importance of Exon 4 sequence. Analysis of *ABI1* mRNA expression during PCa progression revealed differential splicing of the *ABI1* gene involving ABI1-HHR, with Exon 4 expression associated with high grade and stage of PCa tumors and lower survival. Exon 8 and Exon 10 did not show this association (**Fig 5D-F**). Therefore, we can conclude that ABI1 transcripts containing Exon 4 that carry DNA binding property associated with poor prognosis are selected during PCa progression.

**Figure 5.**
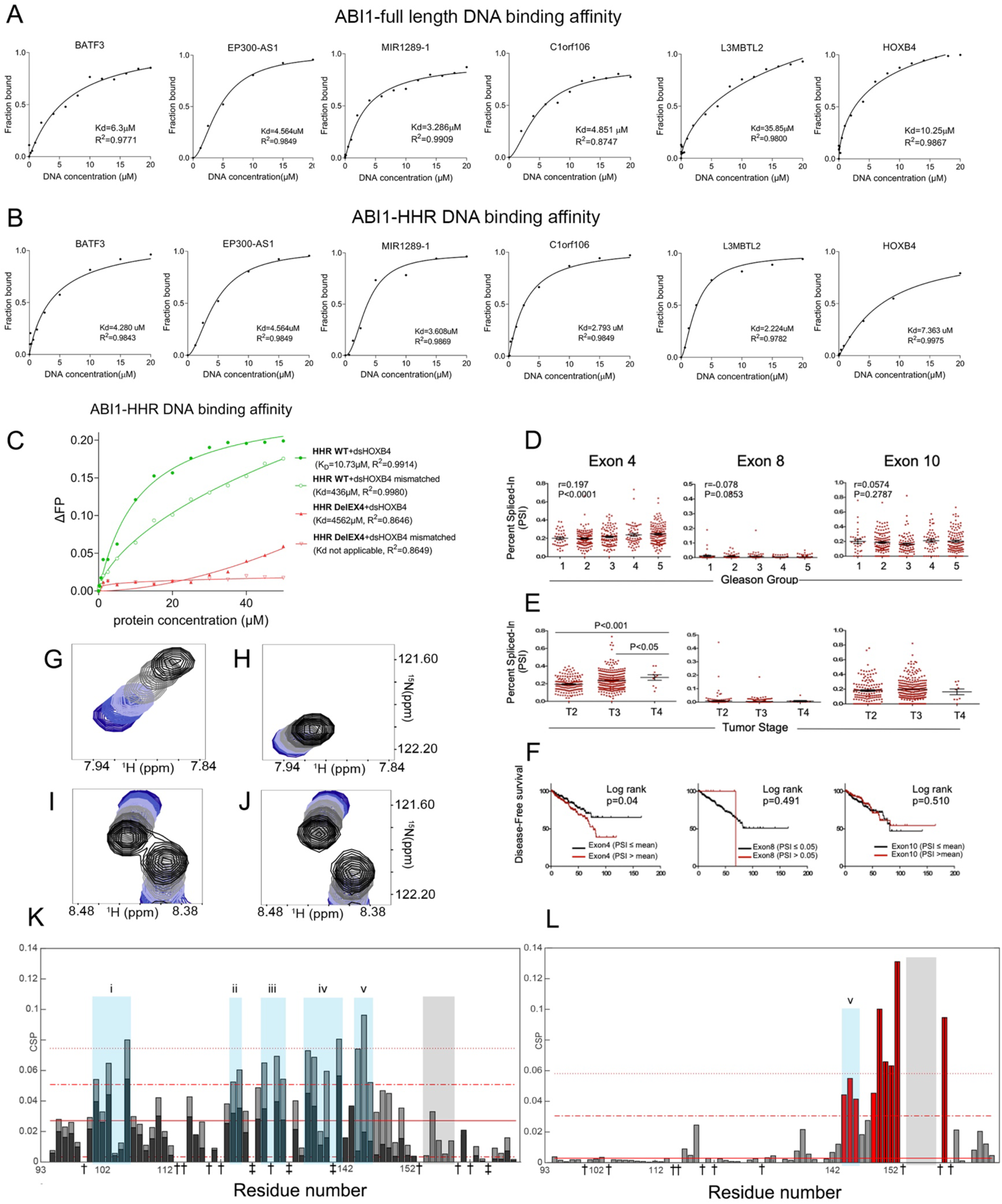
Molecular mechanism of ABI1-HHR DNA binding and its role in PCa tumor progression. **A-B**. Recombinant ABI1 (full-length, **A** or HHR domain, **B**) binding to selected double-stranded (ds) DNA probes. Please note similar affinities of ABI1-HHR and the full-length ABI1; genes symbols are indicated above graphs. **C**, ABI1-HHR binding affinities to HOXB4 dsDNA probe (dsHOXB4) or its mismatched probe (dsHOXB4 mismatched, with 2 non-paring DNA resides in the center); affinities are listed on the right. HHR-WT and HHR-DelEx4, represent wild-type ABI1-HHR or its Exon 4 deleted version, respectively. **D-E.** RNA-seq and the corresponding clinical data of PCa patients from the TCGA cohort (https://www.cancer.gov/tcga) were analyzed for expression of ABI1 Exons 4, 8, and 10; percent of spliced-in (PSI) of ABI1 Exon 4, 8, or 10 are indicated. **D**, Analysis of ABI1 Exon expression within Gleason groups, or within tumor stages (**E**). **F.** Kaplan-Meier curves plot indicates disease-free survival of PCa patients by exon inclusion. ABI1 Exon 4 expression demonstrates lower survival of PCa patients. PCa patients were divided into low- and high-groups according to the PSI level. for each ABI1 Exon separately. **G-J.** Changes in chemical shifts of residues V145 (**G-H**) and K102 (**I-J**) in ABI1-HHR (**G and I**) and ABI1-HHR-DelEx4 (**H and J**) upon gradual addition of HOXB4. Dark blue denotes the free form and black is the bound form, with a color gradient going through grey as the concentration of HOXB4 increases. **K.** Comparison of Chemical Shift Perturbations (CSPs) of ABI1-HHR (grey) and ABI1-HHR-DelEx4 (black) in presence of HOXB4 at stoichiometry 1:3.75. Blue rectangles define regions encompassing groups of residues with CSPs higher than one standard deviation from the median CSP in ABI1-HHR. **L.** CSPs reporting on the deletion of residues 155-159 in ABI1-HHR-DelEx4. Red bars identify residues showing CSPs one standard deviation above the median value. Significant CSPs away from the deletion site reveal allosteric responses. Light grey rectangles in (**K**) and (**L**) highlight residues deleted in ABI1-HHR-DelEx4 (155-159). (‡) identify prolines and (†) identify residues whose peaks could not be centered due to overlap in at least one spectrum required for analysis.

### NMR revealed the molecular determinants for the affinity between DNA and ABI1-HHR

Because deleting Exon 4 seemed to reduce the affinity of ABI1-HHR for HOXB4 and this deletion plays a key role in PCa progression, we used Nuclear Magnetic Resonance (NMR) to identify the molecular determinants for this behavior. NMR reports on changes in the chemical environment of nuclei during binding events making it an ideal tool to probe the interaction with disordered proteins such as ABI1. ABI1-HHR was titrated with HOXB4, and chemical shift perturbation (CSP) analysis revealed five distinct regions (i-v) of ABI1-HHR involved in DNA binding (**Fig. 5K**). The amide signals of the bound form do not display increased dispersion even when approaching saturation, suggesting that the protein does not fold upon binding (**Supplementary Figure S6**) in contrast to many interactions involving IDRs [30]. Instead, the ABI-HHR:DNA interaction appears to occur through a “fuzzy complex” [31], whereby multiple weak, but specific interactions lead to DNA recognition. Most notably, residues in the deleted region representing Exon 4 do not interact with DNA (**Fig. 5K**) and cannot account for a change in affinity upon deletion. To identify the molecular determinants for the change of affinity of ABI1 towards HOXB4 upon deletion of the five residues 155-159 of Exon 4, we titrated ABI1-HHR-DelEx4 with HOXB4 and observed smaller changes in chemical shifts when compared to ABI1-HHR-WT, reflecting a weaker affinity for DNA (**Fig. 5I-J, K**), in agreement with **Fig. 5C**. However, residues in region v and residue 139 in region iv displayed markedly smaller CSPs, indicating that they no longer interact with DNA (**Fig. G-H, K**). To gain insights on the molecular mechanism for this change in affinity, we compared the NMR spectra of apo states of ABI1-HHR and ABI1-HHR-DelEx4. CSP analysis revealed that this deletion not only affects the molecular environment of residues at the deletion site but also alters the molecular environments of residues in binding regions v and iv (**Fig. 5L**). Because residues 155-159 of the deleted region do not interact with HOXB4 in ABI1-HHR, we conclude with confidence that the deletion affects binding through an allosteric mechanism, wherein changes in conformational fluctuations alter residues in regions iv and v such that they can no longer interact with HOXB4.

## DISCUSSION

Here, we identify a novel genomic function of ABI1 in regulating transcription. ABI1 binds DNA using a combination of weak, yet well-defined interactions. We also demonstrate that ABI1 localizes to the nucleus with AR and modifies its transcriptional activity. Parallel changes of ABI1 and AR during PCa progression and treatment indicate that these proteins co-function together in tumor cells. ABI1 levels intimately regulate prostate tissue phenotype, hence, we propose that the ABI1 and AR axis represents a novel mechanism of tumor plasticity in PCa.

ABI1 recognizes DNA through an unusual mechanism that is increasingly reported for disordered proteins, notably for transcription regulation. Protein-nucleic acids interactions have traditionally been thought to be mediated by direct, static contacts between specific protein residues and specific sites on the nucleic acid bases and/or phosphate backbone. While IDRs may undergo disorder-to-order transition when interacting with their binding partners (i.e., fold upon binding), some IDRs with tight binding affinities remain mostly flexible even in their bound states [32, 33]. ABI1 operates through such a mechanism as evidenced by our NMR studies. As pointed out by Fuxreiter *et al* [9] transient, dynamic interactions within the nucleic acid binding interface, as well as flexible regions outside the binding interface can significantly modulate the sequence specificity and/or binding affinity via the formation of fuzzy complexes rather than the adoption of well-defined static structures in the bound state. These fuzzy complexes result in a dynamic DNA readout mechanism, where multiple distinct regions contribute to the conformational ensemble of the complex and the flexibility required to mediate binding. Such disordered motifs are crucially important for transcriptional regulation via protein:protein and protein:nucleic acid interactions, as well as post-translational modifications and/or alternative splicing [34]. ABI1 is involved in all these processes as discussed above, ABI1 PTMS have been reported [7, 8], and hence adopts such a disordered scaffold and binding strategy. As observed for many protein complexes involving IDRs, this lack of stability enhances binding by reducing the loss of conformational entropy upon binding. ABI1 recognizes multiple DNA sequences, and such a binding mechanism may provide versatile target recognition while ensuring functional affinities. Hence, ABI1 may provide fine tuning to transcriptional regulation depending on its binding partner. The DNA sequence sites we have identified will provide a basis to alter binding between ABI1 and its partners, whether through mutagenesis or drug design.

Direct DNA binding of ABI1 indicates that ABI1 may function as a transcriptional factor alone or in collaboration with another transcription factor, such as demonstrated here for AR. ABI1 binds AR in the nucleus and modifies its transcriptional activity, as reflected by changes in AR-regulated gene expression and chromatin occupancy. ABI1 regulates chromatin remodeling proteins and several transcription factor families such as zinc finger binding proteins, homeobox protein, and forkhead proteins, some of which have been implicated in PCa, KLF5 [35], or in multiple cancers such as FoxP2/3 [36]. PLK2 is negatively regulated by ABI1 which suppresses EMT, while it promotes MLKL a key regulator of necroptosis [37] that has been implicated in tumor progression and drug response in PCa [38]. Notably, ABI1 regulates transcription of cell-adhesion genes thus suggesting a positive feedback mechanism for its role in actin cytoskeleton and cell motility. ABI1 and AR transcriptional axis drives a specific gene expression program that can lead to phenotypic changes during PCa progression. Critically, AR acts on ABI1 gene to control the phenotype of prostate epithelium in an androgen dependent manner. AR promotes cell-cell adhesion homeostasis through ABI1, which can be dysregulated after significant changes in AR expression. High levels of AR/ABI1 in CRPC and low in NEPC suggests a phenotypic switch during PCa progression from AR-dependent to AR low/negative state. Moreover, ABI1 is also sensitive to clinical treatments targeting the AR pathway as indicated in by results from NHT samples. Hence, AR-ABI1 axis represents novel mechanism of tumor plasticity in PCa.

Binding data indicate that the critical region responsible for ABI1-DNA binding encompasses the homeobox homologous region termed HHR, residues 93-170 of ABI1. Evolutionary analysis indicates that this region is unique to the ABI protein family in worms, flies, bony fishes, vertebrates, and does not have predictable structural features characteristic of homeobox family proteins (**Supplementary Figure 5)**. Our clinical data indicates the important role of ABI1 Exon 4, coding for residues 155-159, as its expression is selected in high-grade and advanced stage PCa; critically Exon 4 expression is associated with poor survival. We show that deletion of Exon 4 dramatically reduces ABI’s affinity for DNA. NMR spectroscopy analysis indicates that, while not directly contacting DNA, the five residues of Exon 4 affect DNA binding via an allosteric mechanism.

In conclusion, we identified the novel function of ABI1 in DNA binding and transcriptional regulation. ABI1 connects transcriptional activity to cell signaling and actin cytoskeleton, thus exerting cellular and phenotypic plasticity of PCa. Considering widespread expression of *ABI1* in many tissues, AR might be just one example of a transcription factor co-regulated by ABI1.

## Acknowledgments

We thank Dr. P. Kane, J. Ridilla, J. Ross for advice, Dr. A. Perl (Upstate Medical Univ.) for critical reading of the manuscript.

## Author contribution

Concept, direction, overall interpretation-LK; AR-ABI1 binding and localization – BAP, AB; AR-ChIP - KK, AU; ABI1 CHIP – BAP, HU, AU, MN; NMR-DPF and NA designed, performed, and analyzed all experiments; KAM implemented and optimized NMR pulse sequences used for resonance assignments. Protein purification and LLPS – BAP, XL, AK, AB; ABI1-DNA binding-XL, AB; ABI1 splicing-YL, XD; PLA - FZ, SHYK; Histology/TMA-LF, HZO; LuCaP TMA – provided by EC; Clinical Interpretation-MG, GB, MAO, LK. Writing-BAP, NA, MAO, AB, DF, AU, LK. All authors reviewed the manuscript.

**The authors declare no competing interests.**

## Ethics statement

Animal experiments were performed under approved under IACUC protocol No. 375 by the Institutional Animal Care and Use Committee of the State University of New York, Upstate Medical University, Syracuse, NY USA.

## Data access statement

NMR resonance assignments have been deposited in the Biological Magnetic Resonance Data Bank (BMRB) under #51936 for ABI1-HHR, and #51954 for ABI1-HHR-delEx4. ChiPseq and RNAseq will be deposited to public databases for general availability. All primary western blotting data and Excel and/or GraphPad files representing data analysis will be uploaded upon editor request.

## MATERIALS AND METHODS

### Generating ABI1 CRISPR KO and rescue cell lines

The target sequence identification for designing gRNA was performed using the online software <u>crispr.mit.edu</u>. The gRNA sequence CTAGAGGAGGAGATCCCGTC (TGG) in exon 1 of ABI1 was cloned into a pCas-Guide-EF1a-GFP vector (cat. #: <u>GE100018</u>) from Origene Technologies (Rockville, MD). PC3 and LNCaP cells were transfected with the CRISPR plasmid using Lipofectamine 3000 transfection reagent (Thermo Fisher Scientific). Then, the cells were trypsinized and sorted via fluorescence-activated cell sorting (FACS) to obtain GFP-positive cells using a Becton Dickinson FACS Aria III Cell Sorter. Clonal selection of single GFP-positive cells yielded several surviving clones, which were screened for ABI1 protein levels via western blotting, followed by sequence verification by sequencing with ABI1 Exon1 specific primers as described [39]. ABI1 Isoform 2, Isoform 2 (W485N), Isoform 2 (ΔSH3 domain), and Isoform 3 constructs were cloned into a pMSCV puro backbone (Addgene) and packaged into retrovirus using Phoenix-AMPHO producer cells (ATCC). LNCaP ABI1 KO clone 1.5 or PC3 ABI1 KO clone 1.1.1 cells were transduced for two rounds with the retrovirus produced, and the resistant pool was selected with puromycin for use as “rescue” cell lines. Transient transfection of ABI1 KO cells were performed using the TransIT®-2020 (cat. #: MIR 5450) transfection reagent according to manufacturer protocol. Transfection incubation period ranged from 48-72h to obtain maximum plasmid incorporation into cells.

### Cell lines and cell treatments

Authenticated LNCaP, PC3, 22Rv1 (ATCC, Manassas, VA), and LAPC4 cell line (obtained from Dr. Martin Gleave) were maintained in standard RPMI culture media supplemented with 10% fetal bovine serum (HyClone) and the appropriate components and rescue cell lines were supplemented with .25ug/mL puromycin. VCaP cell line (ATCC, Manassas, VA) were maintained in standard DMEM culture media supplemented with 10% fetal bovine serum (HyClone) and the appropriate components. LAPC-4 plates were coated with 5ug/mL poly-L-lysine to help promote adherence to the plate. Cells were maintained in standard phenol-free RPMI 10% charcoal-stripped serum for 3-4 days followed by the addition of Vehicle control (DMSO, androgen deprived media*ADM), 1nM R1881(Sigma, R0908) or 5 μM Enzalutamide (MDV 3100, Selleck Chemicals, S1250) for 5, 16, 24, 48, and 72h. Treatments for live cell assays were performed in CSS unless otherwise noted.

### DNA cloning and constructs

We used the recombinant ABI1 isoform 2 (ABI1-Iso2) [21] [5] for all rescue experiments. To generate SH3 domain mutation, we performed site-directed mutagenesis using Q5 site-directed mutagenesis kit (New England Biolabs Inc.) in the pMSCV-Abi1 Isoform 2 DNA plasmid and point mutated tryptophan at amino acid 485 in the SH3 domain to glutamine. The forward sequence was 5’-GAATGATGATGGCAATTATGAAGGAGTCTGC - 3’ and reverse sequence 5’-GCAGACTCCTTCATAATTGCCATCATCATTC - 3’ synthesized by Integrated DNA Technologies (IDT). To produce the ABI1 Isoform 2 deleted SH3 domain we performed Q5 site-directed mutagenesis (New England Biolabs Inc.) with forward primer 5’-TAGCTCGAGGTTAACGAATTC - 3’ and reverse primer 5’-TTTCTCAATATAATTCTTGGGG - 3’ synthesized by IDT and then verified the sequence (Genewiz). We used the same method to make AR N terminal deletion with forward primer 5’-CCACCCCAGAAGACCTGC-3’ and reverse primer 5’-CCCTAACTGCACTTCTCCAG-3’ synthesized by IDT. The sequence of plasmids were verified by sanger sequencing (Genewiz). For AR we used pEGFP-AR-FL-YFP, pEFGP-ARV7-YFP plasmids in which GFP was replaced it with YFP [40].

### Immunofluorescence and imaging

Ibidi µ-slide 8 well (80827) or #1.5 glass coverslips were coated with collagen IV (Sigma, C5533) and left overnight at 4C. Ibidi wells or coverslips were rinsed with PBS the next day before adding cells. Cells were trypsinized, spun down at 250xg for 4 minutes and resuspended and plated at a concentration of 1×10^3^ cells per well, and maintained for 48h before staining. Cells were then washed with CaCl, MgCl PBS, and fixed with 2-4% paraformaldehyde for 5 minutes at room temperature. Cells were then washed with PBS and quenched with 50mM NH4Cl in PBS for 15 minutes, washed with PBS, blocked with PBSAT (1X PBS, 1% BSA, 0.5% triton-X) for 30 minutes, and then washed with PBS again. Cells were incubated with primary antibody diluted in PBSAT (E-Cadherin, 1:100, Santa Cruz, H-110) overnight at 4C. The next day primary antibody solution was removed from the cells, then the cells were washed with PBS and incubated in secondary antibody diluted in PBSAT (Alexa Fluor donkey anti–rabbit 488, 1:1000, Life Technologies; A21206) for 45 minutes at room temperature. Cells were washed with PBSAT, and then incubated with nuclear counterstain (DAPI, 4,6-diamidino-2-phenylindole, 1:10000, Thermo Fisher, 62248) for 10 minutes. Cells were then washed with PBS and kept in DABCO (1,4-diazabicyclo[2.2.2]octane) antifade reagent (200 mM) for imaging or rinsed with Milli-Q water and mounted on glass slides using Prolong Diamond with 4,6-diamidino-2-phenylindole (DAPI) mounting media and cured in the dark covered for 24 hours before imaging. *Image analysis*: post-image processing was performed using imagJ2. Quantification of E-cadherin immunofluorescence intensity at cell-cell junctions was performed in a minimum of 50 junctions and 5 fields of view in 3 biological replicates, from every condition using line intensity scan mean fluorescence intensity measurements. Images were taken using a epifluorescence spinning disc confocal, 40X water objective. The minimum and maximum threshold were set equally for each image prior to fluorescence measurement. Quantification of total mean fluorescence nuclear intensity of AR was taken from at least 50 cells in 5 different fields of view for 3 biological replicates. Min and Max threshold was set before taking mean intensity measurements to ensure equal intensity values across cell lines; statistical tests performed were One-way ANOVA followed by student’s t-test to indicate statistical differences across cell lines and normalized to either vehicle control or LNCaP ABI1 KO.

### Subcellular fractionation

To isolate subcellular fractions, we washed the cells with ice-cold PBS and then collected the cell pellet at 4C. We then lysed cells and added NE-PER reagents from ThermoFisher (Cat. No.78833). Subsequently, we followed the kit protocol. Protein expression was quantified from densitometry measurements and normalized to fraction specific loading control either cytoplasmic fraction, GAPDH or nuclear fraction, PARP after subtraction of background fluorescence. For each cell line we calculated percentage of nuclear accumulation using the following equation: % Nuclear accumulation = [Nuclear Fraction/(Cytoplasmic fraction + Nuclear fraction)] * 100. Statistical significance was calculated using a One-way ANOVA followed by a student’s t-test between cell lines and calculated from at least 3 biological replicates.

### Immunoprecipitation

Cells had media aspirated and then were immediately washed with cold CaCl/MgCl PBS and lysed on the plate using Pierce IP lysis buffer with protease and phosphatase inhibitors. The resulting lysate was immunoprecipitated using an anti-Abi1 antibody (clone 1B9, MBL) at an antibody:lysate ratio of 1:200 or with anti-IgG1 isotype control (clone G3A1, CST) with the same concentration as an IgG positive control. Immunoprecipitation was performed using Dynabeads™ Protein G (Invitrogen 10004D) according to the manufacturer’s instructions, except the incubation was performed overnight at 4 °C. Samples were washed 3 times with ice-cold PBS and boiled at 90 °C in 2x Laemnli sample buffer (Bio-Rad 1610737) for 5 minutes the following day for elution.

### Immunoblotting

Whole-cell extracts were prepared by lysing cells in 50 mM Tris-Cl pH 7.5, 10 mM MgCl_2_, 0.5 M NaCl, 2% (v/v) NP-40, 0.1% (v/v) benzonase (Sigma-Aldrich E1014), 1% (v/v) Halt^TM^ Protease and Phosphatase Inhibitor Cocktail (Thermo Scientific 78440)Total protein concentrations were determined using Pierce 660 nm (Thermo Scientific 22660) protein assays. Electrophoresis and blotting of protein extracts were performed using mini-PROTEAN tetra handcast and trans-blot turbo transfer systems (Bio-Rad, Hercules, CA).

The primary antibodies used and their working dilutions are listed below. The secondary antibodies (Thermo Scientific) were donkey anti-goat, goat anti-mouse, or goat anti-rabbit IgG HRP conjugates. Blots were imaged using supersignal west pico, pico-plus, or femto chemiluminescent substrate (Thermo Scientific) on a PXi touch imaging system (Syngene, Frederick, MD), and the signal was quantified using GeneTools software (Syngene, Frederick, MD). The data were analyzed using one-way ANOVA and Student’s t-test.

#### Primary antibody used in the study and their dilutions

anti-Abi1 (MBL International, clone 1B9): D147-3 lot# 034, 1:1000

rabbit anti-Abi1 (Cell signaling, D3G6C): 39444 lot# 1, 1:1000

rabbit anti-AR (Santa Cruz, N20): sc-816 lot# N/A, 1:1000

mouse anti-AR (Abcam, 441): ab218431 lot# N/A, 1:1000

rabbit anti-ARV7 (RevMab, clone RM7): 31-1109-00 lot# 091013, 1:1000

rabbit anti-WAVE2 (Santa Cruz, H-110): sc-33548 lot# K0216, 1:1000

rabbit anti-GAPDH (Sigma G9545): G9545 lot# 106M485IV, 1:1000

mouse anti-GAPDH (Cell signaling D4C6R): 97166 lot# N/A, 1:1000

mouse anti-β-actin (Sigma-Aldrich, clone AC-15): A1978, 1:10,000

rabbit anti-E-cadherin (Santa Cruz 7870): sc-7870 lot# N/A, 1:1000

rabbit anti-PARP (Cell Signaling 46D11): 9532 lot# 7, 1:1000

rabbit anti-Stat3 (Cell Signaling 4904): 4904 lot# 7, 1:1000

#### Secondary antibodies used in the study and their dilutions

Donkey anti-Goat IgG (H+L) Secondary Antibody, HRP: A15999 lot# oL1791441c, 1:10,000 Goat anti-Mouse IgG (H+L) Secondary Antibody: PA1-28555 lot# YC3858642E, 1:10,000 goat anti-rabbit IgG HRP: A16096 lot# 96-8-061422, 1:10,000

### Proximity ligation assays

Proximity ligation assay was performed as descried before (Kretschmer A, 2019). The protein-protein interaction was investigated using a commercially available PLA kit (Duolink; Sigma-Aldrich) following the manufacturer’s protocol. Briefly, cells were stained with a pair of primary antibodies at 4 °C for overnight and then labeled with probes linked with complementary DNA (cDNA) at 37 °C for 1 hour. cDNAs were ligated at 37 °C for 40 minutes and the product was amplified at 37 °C for 100 minutes. The signal was inspected using the Olympus FV3000RS confocal microscope with a 60x UPLAPO oil objective (Olympus Canada Inc., Richmond Hill, Ontario, Canada; Manufactured by Olympus in Tokyo, Japan.). The images were taken using the Z-stack model with the confocal pinhole set to “automatic” with the FC31S-SW software. For presentation purpose, images were exported as bitmap (BMP) files [41].

### Recombinant Protein Expression

All His-SUMO-AR and ABI1 constructs were transformed and expressed in BL21(DE3) codon plus *E. coli* cells using standard molecular biology techniques. Briefly, plasmid transformations were performed by incubating cells on ice for ∼30 minutes, followed by heat shock at 42°C for 30 seconds and subsequent icing for 1 minute. LB broth media was added to cells to allow recovery at 37 °C with shaking for 1 hour. Cells were spread/grown on LB-agar plates supplemented with the selection antibiotics chloramphenicol (CP) and kanamycin (kan), and then were incubated overnight at 37 °C. The following morning, colonies from the plates were picked and used to inoculate a 5 ml LB media containing CP/Kan and grown at 37°C for 8 hours with shaking. Cells cultures were then scaled up to a 250 mL flask containing 50 mL LB media with CP/kan and 0.5% glucose, and then allowed to grow overnight at 37°C with shaking. The following day, 10-20 mL of the overnight cell culture was then transferred into 2.8 L fernbach flasks containing 1 L of LB media supplemented appropriate antibiotics (CP/Kan) and allowed to grow at 37°C until an OD _600 nm_ between 0.6. -1.0 was reached. Then, protein expression was induced with 1 mM IPTG and grown overnight at 16°C with shaking. The following morning, the cells were harvested via centrifugation at 3500 rmp at 4°C. Cell pellets were then resuspended in NiA-Guad buffer (6M Guanidium-HCl, 300 mM NaCl, 50 mM Na_2_PO_4_, 5 mM betamercapto-ethanol, 20 mM Imidazole pH 7.5), and stored at -20°C until needed for purification.

### Protein Purification

Cell pellets were thawed and sonicated with pulsing of 2 secs on and 4 secs off for 5 minutes to lyse the cells, followed by centrifugation to remove any insoluble cell debris. Supernatant, which contains the protein of interest, was then added to a column containing Nickel-NTA beads, pre-equilibrated with NiA-Guad buffer. To allow the His-SUMO-tagged protein of interest to bind, the Ni^2+^ column was incubated at 4°C for 45 minutes with gentle agitation. After incubation, unbound protein was then collected in the flow-through fraction and stored for later purification. For subsequent Ni^2+^column protein purification of bound protein, the column was washed 3 times with ∼30 mL of NiA-Guad buffer and then 3 times with NiA-Arg-Gly buffer (300 mM NaCl, 200 mM Arg-HCl, 10% Glycerol, 50 mM Na_2_PO_4_, 5 mM betamercapto-ethanol, 20 mM Imidazole pH 7.5). Protein was eluted off Ni^2+^ column beads using NiB-Guad buffer (i.e. NiA-Guad supplemented with 500 mM Imidazole) in 8 elution fractions of ∼10 mLs each. SDS-PAGE gel electrophoresis was used to identify the fractions containing the protein of interest and their purity. We then combined elution fractions that had the most abundant and pure protein, concentrated them and then load them them on size-exclusion 26/60 gel filtration column (Cytiva) for further purification in NiA-Guad buffer. Again SDS-PAGE gel were used to identify fractions containing pure protein, which were then combined, and then concentrated to around 7-10mg/ml using centrifugation at 4°C. Some of the proteins of interest were labelled with Alexa Fluor™ 488 C_5_ Maleimide (Thermofisher) according to the manufacturer’s instructions. We dialyze all our proteins twice to perform buffer exchange overnight at 4°C into 4 L buffer of interest prior to performing our phase separation or binding assays.

### DIC LLPS and turbidity assay

All buffers used are passed through 0.22µm filters to be used as buffer control condition. We performed turbidity assays to monitor LLPS using a Cary 3500 UV-Vis double-beam, multicell spectrophotometer system (Agilent) with varying concentrations of protein, NaCl and temperature. Cuvettes were pre-cooled in instrument before each experiment. Fluorescently-labelled samples were used to perform Differential Interference Contrast (DIC) microscopy using an Axio-Zeiss upright microscope with 100x oil objective or using FITC channel for ABI1-488.

### Statistical test

All statistical tests were run with student’s t-test or ANOVA from 3 biological replicates. Turbidity assays using cloud point temperatures were calculated with 4PL regression analysis as this was the best fit for the data.

### ABI1 FL/HHR:DNA binding assays by fluorescence quenching

Binding buffers, Low salt: 1X PBS (pH=7) supplemented with 5 μM betamercapto-ethanol, High salt: 1X PBS (pH=7) supplemented with 500 mM NaCl and 5 μM betamercapto-ethanol. Double-stranded DNA (dsDNA) annealing, and preparation was performed with pre-mix 1 mM single-stranded DNA (ssDNA) with its reverse complement strand in appropriate buffer (low salt binding: 1XPBS; high salt binding: 1XPBS with 500 mM NaCl) to make annealing reaction containing 100 μM of the complementary ssDNA that are heated to 98°C and cooled slowly to 4°C in a thermal cycler. After annealing reaction, we perform electrophoresis on a 4% agarose gel to confirm annealing efficiency using our ssDNA as controls. For fluorescent labeling our DNA oligos of interest, we use a 5’-C6 amino modified dsDNA to covalently link a Alexa Fluor™ 488 NHS dye according to the manufacturer’s instructions.

To detect protein intrinsic fluorescence quenching effect upon adding dsDNA, we use “buffer+30 μM dsDNA” as negative control (baseline fluorescence) and buffer + 3 μL HHRWT as maximum fluorescence reading. Perform the fluorescence spectrum reading using Spectra Max® i3 plate reader from Molecular Device. Set excitation wavelength at 250 nm and emission wavelength range from 280 nm to 400 nm, 5 nm per fluorescence read. To detect fluorescence quenching from Alexa dye labelled dsDNA upon adding proteins, use reading from “Buffer + 30 μM HHR proteins” as negative control (base line fluorescence). Use buffer + 1 μL Alexa dsDNA as maximum fluorescence reading. Preparing titrated HHR proteins with buffer in the following wells and add 1 μL of Alexa dsDNA into each well. Perform the fluorescence spectrum reading using Spectra Max® i3 plate reader from Molecular Device. Set excitation wavelength at 360 nm and emission wavelength range from 400 nm to 650 nm, 5 nm per fluorescence read. Export the raw fluorescence data with peak fluorescence reading for each titrated sample and subtract that value by the reading from negative control well. Set the reading from maximum fluorescence reading well (Buffer +3 μL HHR WT for protein intrinsic fluorescence, Buffer + 1 μL dsDNA for Alexa fluorescence) as 100% and normalized other fluorescence readings from titrated wells to the maximum fluorescence reading well. Convert normalized fluorescence to fraction bound by using the equation: Fraction Bound = 1-Normalized fluorescence. Plot concentration of dsDNA/proteins to fraction bound and fit the scatter dot plot with Nonlinear regression curve fit function with the specific binding with hill slope function in Prism to obtain the dissociation constant (K_d_). To detect changes of fluorescence polarization from Alexa dye labelled dsDNA upon adding proteins, use reading from buffer + 1 μL Alexa dsDNA as baseline fluorescence polarization. Preparing titrated HHR proteins with buffer in the following wells and add 1 μL of Alexa dsDNA into each well. Perform the fluorescence polarization reading using Spectra Max® i3 plate reader from Molecular Device. Set excitation wavelength at 485 nm and emission wavelength at 535 nm. Extract raw fluorescence intensity and calculate the fluorescence polarization (FP) using following equation: FP=(F_parallel_ - F_perpendicular_) / (F_parallel_ + F_perpendicular_), where F_parallel_ = fluorescence intensity parallel to the excitation plane, and F_perpendicular_ = fluorescence intensity perpendicular to the excitation plane. Calculate ΔFP from titrated wells to buffer + 1 μL Alexa dsDNA well by subtracting FP_baseline_ from FP_titrated wells_. Plot concentration of proteins to ΔFP and fit the scatter dot plot with Nonlinear regression curve fit function with the specific binding with hill slope function in Prism to obtain the dissociation constant (K_d_).

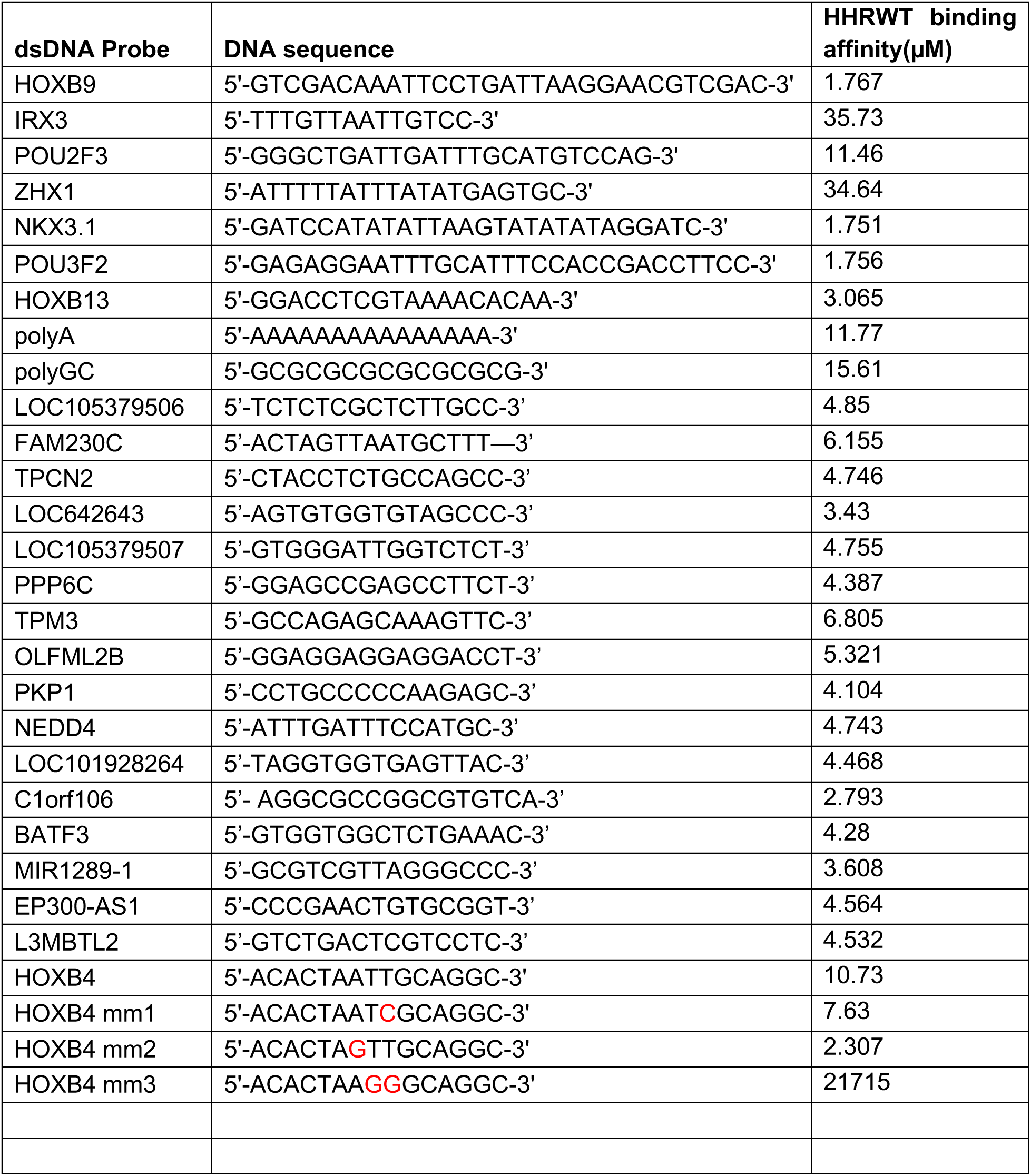

### RNA isolation and Quantitative PCR analysis

Total RNA was extracted from cell lines using the Qiagen shredder and RNAeasy Extraction Kit (Qiagen) and then reverse-transcribed using iScript cDNA synthesis kit (Bio-Rad Laboratories, Inc., Hercules, CA). We then added 10ng template to each qPCR assay with FAM/ZEN/IBFQ detection and performed real-time qPCR amplification using CFX Connect system (Bio-Rad Laboratories, Inc., Hercules, CA). Primetime qPCR assays (IDT) were used to screen for ABI1, AR, KLK3, TMPRSS2, FKBP5, Wasf1, and GAPDH expression across multiple LNCaP sublines with qPCR assays Hs.PT.58.40787474, Hs.PT.56a.14520219, Hs.PT.58.38546086, Hs.PT.58.39738666, Hs.PT.58.39408001, Hs.PT.58.2775672, Hs.PT.39a.22214836 and followed the manufacturer’s instructions. Relative gene expression was calculated using the following _equations: R=2−[(Cq;GOIA)−Cq;REFA)−(Cq;GOIB−Cq;REFB)]= 2−(ΔCq;A−ΔCq;B)=2−ΔΔCq_ [42]_. Statistical tests_ performed were One-way ANOVA followed by student’s t-test to indicate statistical differences across cell lines and each target gene was normalized to loading control GAPDH followed by ABI1-Iso2 Rescue cell line.

### AR ChIP-qPCR and ABI1 qPCR analysis

ABI1 2^nd^ intron / KLK3 AREIII (positive control region) AR ChIP-qPCR with samples. ChIP samples: LNCaP-pcDNA, ARmo and ARhi 2h stimulation of 0, 1 or 100 nM DHT + inputs and IgG = 36 samples in duplicates = 72 + standards in duplicates and NTC = 14. Run with CFX96 Real-Time PCR System (BioRad). SYBR green assay components: Maxima SYBR Green Master Mix(2x)=10 μL, Primer(forward and reverse, 100 μM)=0 or 1 μL, Template DNA=500ng (1 μL), Water= 8 μL; totally assay volume=20 μL. PCR protocol: 95°C for 10 min (1 cycle), 95°C for 15 s (40x cycle), 54 °C for 30 s (40x cycles), 72°C for 30 s (40x cycles), 60-95°C (1 cycle). Sequences of the primers targeting second intron of ABI1: ABI1_intF3-AATCCCAGCTACTCACTCGG; ABI1_intR3-AGTCTCGCTCTGTCACCCA. The data are presented as fold over percentage of input of the ethanol-treated sample. Mean ±s.e.m. of three technical replicates are shown [43, 44]. The pull downs are the same as performed in, Urbanucci, A., et al. (2012) [43].

### RNA sequencing

Total RNA was extracted from cell lines using the Qiagen shredder and RNAeasy Extraction Kit (Qiagen). Statistical and quality control analysis was performed using Partek bioinformatics software. Unaligned reads were trimmed based on quality score(min quality phred score=20); STAR alignment(genome build homo sapiens; aligner index whole genome); quantification to annotation model using refseq annotation (Partek E/M); filtered features by filtering out low expression at gene level, excluding features if maximum <=10; normalized counts for differential expression analysis using, total count=total # reads in data/total # reads in genome and adding .0001=to assist in offset in statistics; performed differential expression analysis using treatment and cell type as attributes to include in statistical test with comparisons between: ISO2/KO/Parental 0nM treatment (DMSO only) vs ISO2/KO/Parental 1nM (R1881) treatment, ISO2 0nM treatment vs ISO2 1nM treatment, KO 0nM treatment vs KO 1nM treatment, parental 0nM treatment v parental 1nM treatment, ISO2 0nM treatment vs KO 0nM treatment, ISO2 1nM treatment vs KO 1nM treatment, Parental 0nM treatment vs KO 0nM treatment, and Parental 1nM treatment vs KO 1nM treatment.

### ChIP sequencing

For ChIP analysis we used anti-Abi1 antibody (MBL International, cat# D147-3, Lot# 034), AR(N20, SCBT, sc-816). Each ChIP reaction was carried out using 40 ug of chromatin sample. ChIP data analysis was generated by ActiveMotif, Inc using the following software: bcl2fastq2 (v2.20): processing of Illumina base-call data and demultiplexing. bwa (v0.7.12): alignment of reads to reference genome. Samtools (v0.1.19): processing of BAM files. BEDtools (v2.25.0): processing of BED files. MACS2 (v2.1.0): peak calling; narrow peaks. SICER (v1.1): peak calling: broad peaks. wigToBigWig (v4): generation of bigWIG files. AR and ABI1 chromatin binding identified using genomic regions containing 1 or multiple overlapping Intervals and the associated FDR or “padj” cutoff for significance FDR < 0.001 based on DESeq2 analysis of 50kb. DESeq2 analysis read counts for all Merged Regions are first obtained from the unnormalized BAM files (without duplicates). The program then normalizes the counts between the samples using the “median of ratios” method, and calculates log2 fold-change (Log2FC), shrunken-Log2FC, p-value, and adjusted p-value (padj, FDR) for each Merged Region.

Sequencing depth and peak calling parameters. The ChIP DNA was processed into a standard Illumina ChIP-Seq library and sequenced to generate >5 million reads. ABI1: Reads (8.6 million) were aligned to the human genome (hg38), and after removal of duplicate and non-uniquely mapped reads, only ∼2.13 million alignments were obtained. AR: Reads (>35 million) were aligned to the human genome (hg38), and after removal of duplicate and non-uniquely mapped reads, only ∼26 million alignments were obtained. A signal map capturing fragment densities along the genome was generated and visualized in the Integrated Genome Browser (IGB). In addition, MACS peak finding was performed to identify the most significant peaks. Data quality: Using our default cutoff of p-value 1e-7, (ABI1) 1,587 peaks & (AR) >41,000 were identified (after ENCODE blacklist filtering).

### Data availability statement

ChiPseq, RNAseq, and NMR data will be deposited to public databases for general availability. All primary western blotting data and Excel and or GraphPad files representing data analysis will be uploaded prior final acceptance/upon editor request.

### Sample size and data reproducibility

Each experiment was performed in 3 biological and 3 technical repeats. Data from all technically well executed experiments were included. Samples were randomized. ABI1 expression in mice and human histopathology samples were blindly evaluated. RNAseq was performed in triplicate sample. ChIP seq was performed in duplicate samples with high correlation, thus justifying sample size.

### Cell lines

Authenticated cell lines were obtained from ATCC. All cell lines tested negative for Mycoplasma. No misidentified cell lines were used in the study.

### Animals

All animal experiments were performed under approved under IACUC protocol No. 375 by the Institutional Animal Care and Use Committee of the State University of New York, Upstate Medical University, Syracuse, NY USA. C57BL/6J Mice were used for the study, only male mice were used to determine expression of Abi1 upon surgical castration for 4 weeks, n=3.

### Prostate cancer cohorts

#### CRPC-NEPC cohort

A TMA containing tissue cores patients who had undergone no treatment, neoadjuvant hormonal therapy (NHT), chemotherapy, or radiotherapy was obtained from the tissue biobank at the Vancouver Prostate Centre. Histopathology of the primary treatment naïve tumors (n=63), NHT-treated tumors under 6 months (n=71), over 6 months (n=67), all NHT treated (n=159), CRPC (n=22), and NEPC-like (n=17) tumors has been previously reported and characterized [45]. *NHT clinical cohort* was previously published in [46].

### Immunohistochemistry (IHC)

Staining of patient tumor sections was completed on a Ventana DISCOVERY Ultra autostainer (Ventana Medical System, Tuscan, Arizona) with an enzyme labeled biotin streptavidin system and solvent resistant DAB Map kit using corresponding primary antibody. Antigen retrieval was performed with Cell Conditioning 1 (CC1) (Ventana) at 95°C for 64 minutes. Slides were incubated with anti-ABI1 (4E2), 1:200 of “anti-Sox9” antibody (AB5535, Millipore Sigma) at room temperature for 2 hours. The UltraMap DAB anti-Rb Detection Kit (Ventana) was then used for detection. All stained slides were digitalized with Aperio AT2 Scanner (Leica Biosystems) at 20X magnification. The images were subsequently stored in the SlidePath digital imaging hub (DIH; Leica Microsystems) of the Vancouver Prostate Centre. The area of interest in the tumor images were delineated by pathologist (Dr. Ladan Fazli) and automated digital image analysis were run for each biomarker using Aperio Positive Pixel Count Algorithm.

Tissue microarray samples and quantification (LuCaP)

TMA data of ABI1 Immunoexpression quantified using 3,3′-Diaminobenzidine Optical Density (DAB OD) with QuPath software of LuCaP Cell lines: LuCaP 35/136/81; LuCaP 35CR/81CR/136CR; LuCaP 77CR/86.2/93/145.1/145.2. Statistical significance was evaluated using one-way ANOVA, followed by t-test, n=3.

### NMR Spectroscopy

Isotopically enriched HHR WT/DelEx4 samples were grown in M9 minimal media containing ^15^NH_4_Cl and ^13^C glucose (Cambridge Isotope Labs) as the sole source of nitrogen and carbon respectively. Protein expression and purification was performed as described above for the unlabeled samples. All NMR spectroscopy data were acquired at 20°C in NMR buffer (150 mM KCl, 20 mM HEPES, 5 mM betamercapto-ethanol, pH 7) on a 600 MHz Bruker Avance III spectrometer equipped with a QCI probe.

### NMR resonance assignment

Unambiguous assignment of ABI1-HHR and ABI1-HHR-DelEx4 was achieved by conducting triple resonance experiments with a 240 μM ^15^N, ^13^C doubly labeled sample of ABI1-HHR (93-170). Eight 3D experiments: 3D HNCO (4 scans, 1024 (^1^H, 10.58 ppm at 4.696 ppm) x 64 (^15^N, 25 ppm at 120.5 ppm) x 75(^13^C, 7 ppm at 173.1 ppm) complex points, 2 h 47 min), 3D HN(CA)CO (16 scans, 1024 (^1^H, 10.58 ppm at 4.696 ppm) x 64 (^15^N, 25 ppm at 120.5 ppm) x 75(^13^C, 7 ppm at 173.1 ppm) complex points, 11 h 14 min), 3D CBCA(CO)NH (16 scans, 1024 (^1^H, 10.58 ppm at 4.696 ppm) x 36 (^15^N, 25 ppm at 120.5 ppm) x 58 (^13^C, 58 ppm at 41 ppm) complex points, 4 h 54 min), 3D HNCA (8 scans, 1024 (^1^H, 10.58 ppm at 4.696 ppm) x (^15^N, 25 ppm at 120.5 ppm) x (^13^C, 24.4 ppm at 49.75 ppm) complex points, 7 h 16 min), 3D HNCACB (16 scans, 1024 (^1^H, 10.58 ppm at 4.696 ppm) x 64 (^15^N, 25 ppm at 120.5 ppm) x 150 (^13^C, 58 ppm at 41 ppm) complex points, 22 h 2 min), 3D (H)N(CA)N(N)H (16 scans, 1024 (^1^H, 10.58 ppm at 4.696 ppm) x 128 (^15^N(t_1_), 25 ppm at 120.5 ppm) x 128 (^15^N(t_2_), 25 ppm at 120.5 ppm) complex points, 10 h 23 min), 3D (HACA)N(CA)CON (24 scans, 1024 (^13^C, 7 ppm at 173.1 ppm) x 90 (^15^N(t_1_), 33.4 ppm at 123.4 ppm) x 90 (^15^N(t_2_), 33.4 ppm at 123.4 ppm) complex points, 1d 22 h 42 min), 3D (HACA)N(CA)NCO (88 scans, 1024 (^13^C, 7 ppm at 173.1 ppm) x 90 (^15^N(t_1_), 33.4 ppm at 123.4 ppm) x 90 (^15^N(t_2_), 33.4 ppm at 123.4 ppm) complex points, 4d 4 h 14 min) were recorded. A 4D HNHsNs Covariance map (Harden & Frueh, 2018) [47] was calculated from HNCO, HN(CA)CO, CBCA(CO)NH, and HNCACB to prevent errors in peak picking and ensure that sequential residues were reliably assigned. All experiments were run with non-uniform sampling using a 10% NUS sampling factor, processed in nmrPipe (Delaglio et al, 1995) [48], and analyzed in CARA methods from Keller, 1996 [49].In the end, all residues of ABI1-HHR could be assigned except for the first two N-terminal residues.

### NMR titrations

^15^N ABI1-HHR and ^15^N ABI1-HHR-DelEx4 were titrated with HOXB4 DNA with stoichiometries 1:0.47, 1:0.94, 1:1.9, 1:3.75, and 1:7.5 and monitored by NMR. ^15^N ABI1-HHR, ^15^N ABI1-HHR-DelEx4, and HOXB4 were buffer exchanged into the same freshly prepared buffer solution. The titrations were performed at constant protein concentration (80 µM) by mixing solutions of free ABI1 with those of ABI1 in presence of DNA at various concentrations, starting with a solution with 7.5 equivalent of DNA (600 µM). 1:375 was obtained by mixing free ABI1 with 1:7.75 in equal amounts, 1:1.9 by mixing free ABI1 with 1:3.75, 1.094 by mixing free ABI1 with 1:1.9, and 1:0.47 by mixing free ABI1 with 1:0.94. All samples contained 10% D_2_O and 200 μM sodium trimethylsilylpropanesulfonate (DSS) chemical shift referencing. All titration points were recorded sequentially in the order mentioned above at a temperature T = 20°C as calibrated with methanol. This data collection scheme ensures that chemical shift perturbations report exclusively on binding and are not tainted by sample degradation. ^15^N-^1^H HSQC experiments were run with 16 scans, 1024 (^1^H, 16 ppm at 4.696 ppm) x 256 (^15^N, 25 ppm at 120.5 ppm) complex points (2 h 45 mins). Chemical Shift Perturbations were calculated with an in-house MATLAB script using the formula from Williamson, 2013 [50]:

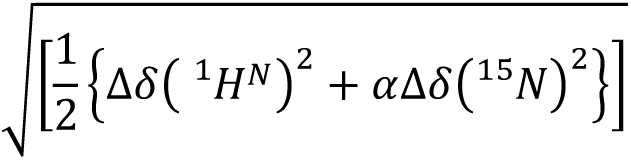

A scaling factor, α = 0.2 was used for ^15^N chemical shifts.

