## Supplemental Figures for "ABI1 regulates transcriptional activity of Androgen Receptor by novel DNA and AR binding mechanism"

**SUPPLEMENTARY FIGURES, Porter, Li, *et al* 2023**

A

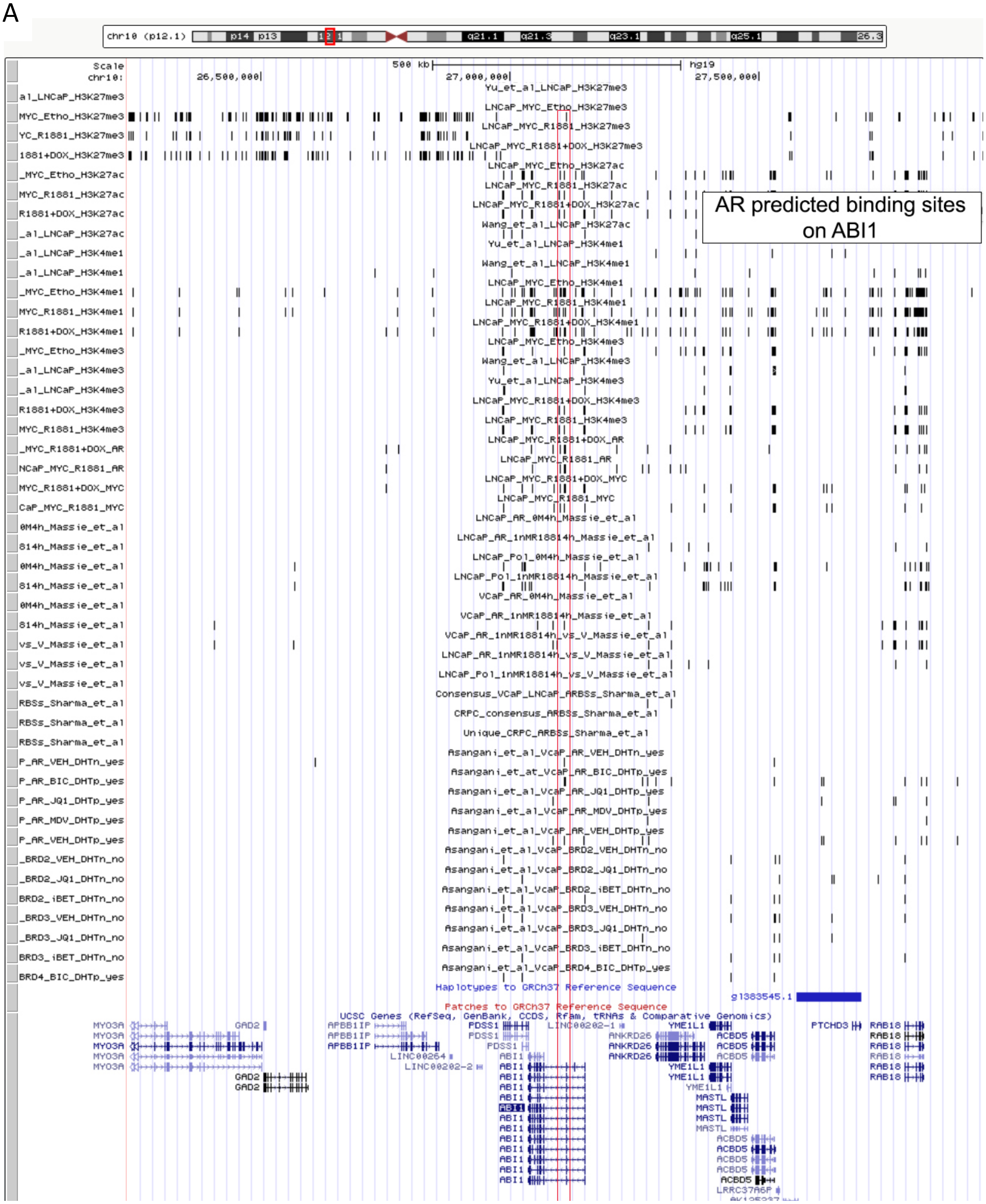

B

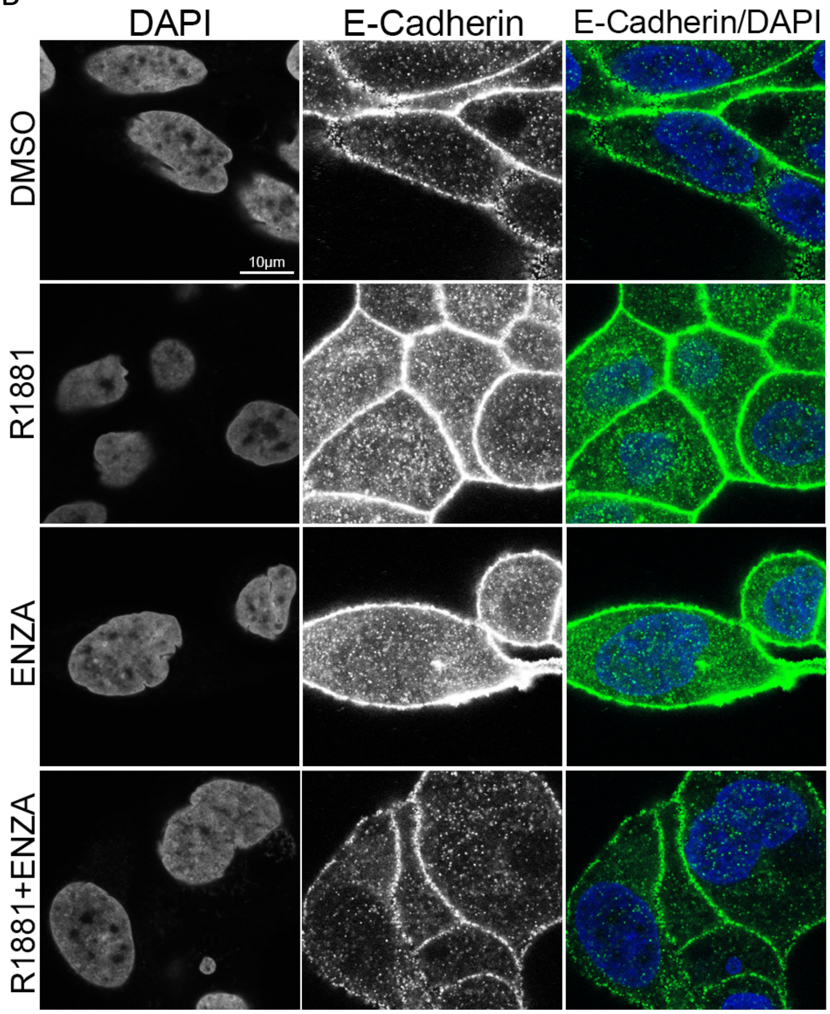


**Supplementary Figure 1. A. AR binds to *ABI1* gene and pharmacological treatments can shift chromatin binding location.** The figure depicts data sets selected for analysis on tracks in UCSC genome browser resources. Black lines indicate binding events at specific locations across chromatin. Red box indicates where AR predicted binding locations on ABI1 is. Treatments that can shift AR associated ABI1 chromatin binding sites include MDV, iBET, JQ1, and Bicalutamide (BIC). AR binning versus MYC is indicated in heading above track. Histone methylation and acetylation events are indicated in the top 18 tracks. Active methylation is associated around the AR predicted binding sites on ABI1 indicating that this binding event is indeed active and can induce gene expression. Data sets include: patient data [1] [2], CRPC samples [3], unique AR binding events [3] (Sharma et al 2013), cell line data VCaP [4] [5] , LNCaP [5], and [6].

**B. Enzalutamide inhibits cell-cell adhesion in LNCaP cells.** Example images demonstrating the effect of anti-AR treatment on downstream function of ABI1 in promoting E-cadherin localization to cell-cell junctions. Stimulation of AR with synthetic androgen (R1881) compared with no ligand control (DMSO) led to increased E-cadherin localization at the cell-cell junction and formation of cohesive cell to cell contacts with neighboring cells. Antagonism of AR with enzalutamide led to decreased cell to cell contacts with neighboring cells indicated by minimal cell junction formation as shown with E-cadherin and/or large gaps between cells (ENZA). Simultaneous ligand stimulation with antagonism of AR (R1881 + ENZA), bottom panel, led to weaker E-cadherin signal intensity at the cell-cell contacts. Quantification of E-cadherin peak intensity at cell-cell junction is presented in main **Figure 2H**.

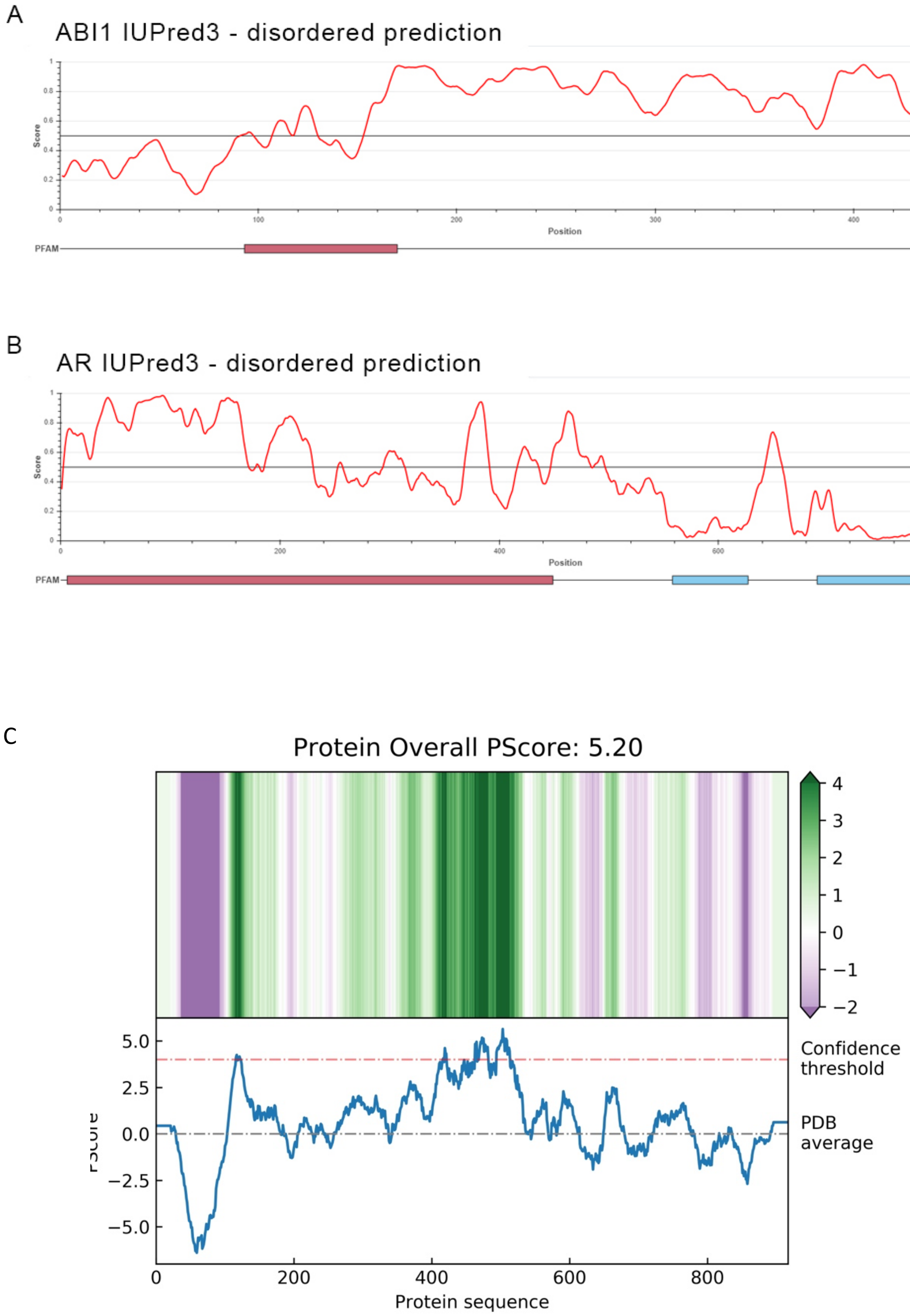
A
ABI1 IUPred3 - disordered prediction
IUPred3 (long)
Score
Position
PFAM
B
AR IUPred3 - disordered prediction
IUPred3 (long)
Score
Position
PFAM
C
Protein Overall PScore: 5.20
Confidence threshold
PDB average
Protein sequence

D

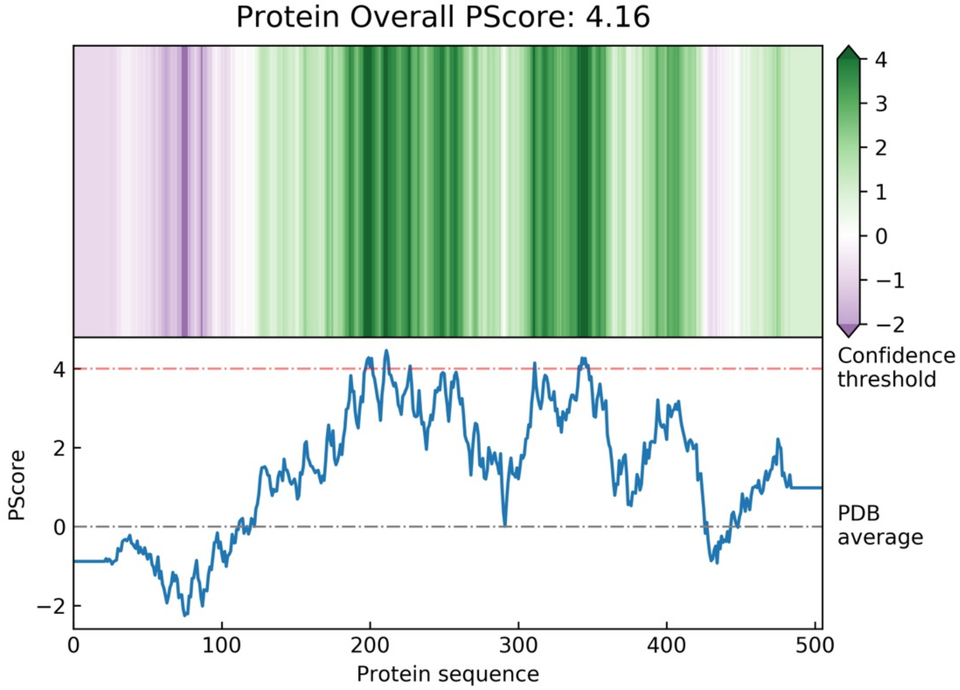


**Supplementary Figure 2. Predicting the ABI1 and AR intrinsically disorder regions and liquid-liquid phase separation. A.** ABI1 disordered prediction using IUPred3 showing that ABI1 is disordered (score > 0.5) through most of the protein except the highly structured C-terminal SH3 domain and a region in the N-IDR. **B.** AR-full length disordered prediction using IUPred3 showing that the N-terminal region up until the ligand binding domain (LBD) is highly disordered. **C & D.** PScore plots for AR and ABI1 respectively showing the likelihood of their IDRs to phase separate based on propensity for long-range planar pi-pi interactions. Both AR and ABI1 have overall scores above the confidence threshold for each protein.

A

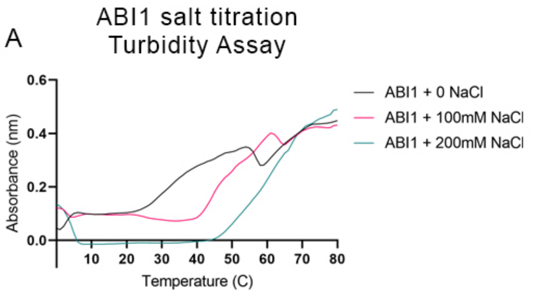


B

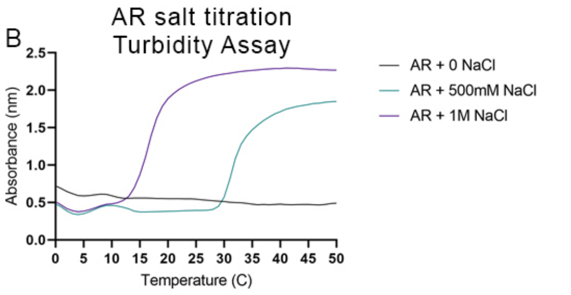


a

| Sample | Inflection Point/Cloud Point Temperature |
|---|---|
| ABI1+0 NaCl | 46.69 |
| ABI1+100mM NaCl | 50.40 |
| ABI1+200mM NaCl | 61.39 |

b

| Sample | Inflection Point/Cloud Point Temperature |
|---|---|
| AR+0 NaCl | 10.54 |
| AR+500mM NaCl | 32.70 |
| AR+1M NaCl | 17.35 |

C

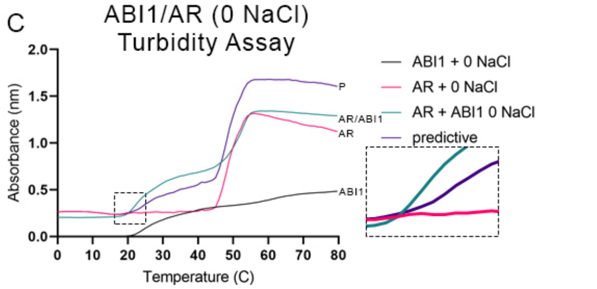


c

| Sample | Inflection Point/Cloud Point Temperature | $\Delta$Temp=Final$_{(AR+ABI1)}$-X$_{(ABI1\ or\ AR)}$ |
|---|---|---|
| AR+0 NaCl | 48.55 | -4.81 |
| ABI1+0 NaCl | 37.60 | +6.13 |
| AR+ABI1+0 NaCl | 43.73 | 0 |

D

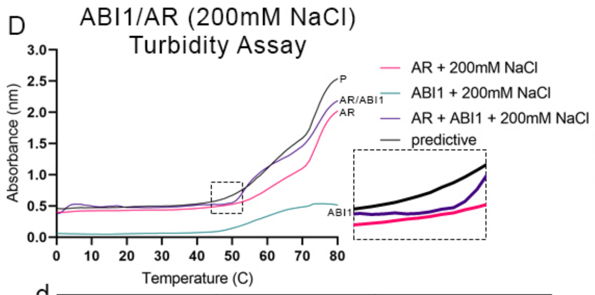


d

| Sample | Inflection Point/Cloud Point Temperature | $\Delta$Temp=Final$_{(AR+ABI1)}$-X$_{(ABI1\ or\ AR)}$ |
|---|---|---|
| ABI1+200mM NaCl | 57.23 | +7.08 |
| AR+200mM NaCl | 68.19 | -3.87 |
| AR+ABI1+200mM NaCl | 64.32 | 0 |

E

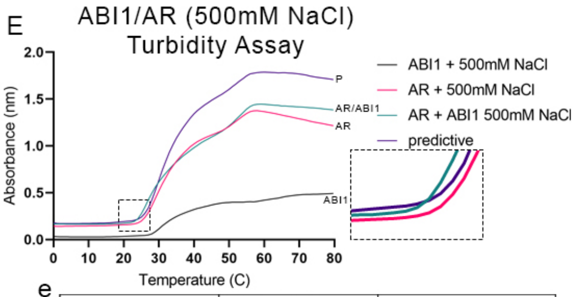


e

| Sample | Inflection Point/Cloud Point Temperature | $\Delta$Temp=Final$_{(AR+ABI1)}$-X$_{(ABI1\ or\ AR)}$ |
|---|---|---|
| AR+500mM NaCl | 33.34 | +1.62 |
| ABI1+500mM NaCl | 35.59 | -0.63 |
| AR+ABI1+500mM NaCl | 34.96 | 0 |

F

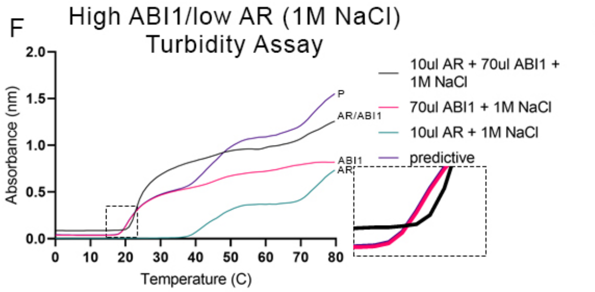


f

| Sample | Inflection Point/Cloud Point Temperature | $\Delta$Temp=Final$_{(AR+ABI1)}$-X$_{(ABI1\ or\ AR)}$ |
|---|---|---|
| 10ul AR+1M NaCl | 90.45 | -61.25 |
| 70ul ABI1+1M NaCl | 30.07 | -0.86 |
| 10ul AR+70ul ABI1+1M NaCl | 29.20 | 0 |

G

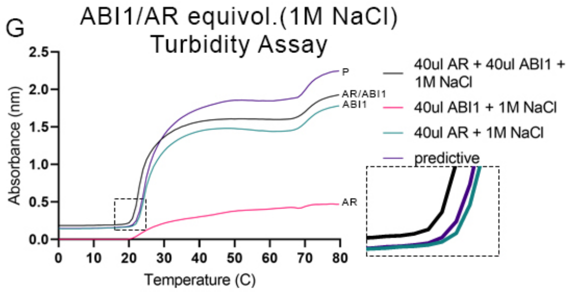


g

| Sample | Inflection Point/Cloud Point Temperature | $\Delta$Temp=Final$_{(AR+ABI1)}$-X$_{(ABI1\ or\ AR)}$ |
|---|---|---|
| 40ul AR+1M NaCl | 26.45 | -1.71 |
| 40ul ABI1+1M NaCl | 33.07 | -8.33 |
| 40ul AR+40ul ABI1+1M NaCl | 24.74 | 0 |

**Supplementary Figure 3. Phase separation of ABI1 and AR. A.** ABI1 salt titration and cloud point temperatures based on turbidity assays suggest that ABI1 undergo LLPS under low salt and low temperature conditions implicating a UCST nature of the protein's cloud point temperature (table a). **B.** AR salt titration and cloud point temperatures based on turbidity assays suggest that AR undergo LLPS under high salt and high temperature conditions implicating a LCST nature of the protein's cloud point temperature (table b). **C.** AR and ABI1 in a 0 mM NaCl condition indicating the combination of AR and ABI1 in the same solution has a shift in cloud point temperatures +/-, ABI1 +6.13C, AR -4.81 (table c). **D.** AR and ABI1 in a 200 mM NaCl condition indicating the combination of AR and ABI1 in the same solution has a shift in cloud point temperatures +/-, ABI1 +7.08C, AR -3.87 (table d). **E.** AR and ABI1 in a 500 mM NaCl condition indicating the combination of AR and ABI1 in the same solution has a shift in cloud point temperatures +/-, ABI1 -0.63, AR +1.67 (table e). **F.** AR and ABI1 in a 1 M NaCl condition with high ABI1 (70 ul) and low AR (10 ul) indicating the combination of AR and ABI1 in the same solution at different volumes has a unique shift in cloud point temperatures +/-, ABI1 -0.86, AR -61.25 (table f). **G.** AR and ABI1 in a 1 M NaCl condition with equivolumetric AR and ABI1 indicating the combination of AR and ABI1 in the same solution at the same volume has a unique shift in cloud point temperatures +/-, ABI1 -8.33, AR +1.71 (table g).

**Predictive curve values indicate sum of individual protein values from turbidity measurements which is different from turbidity measurements taken when they are combined into one solution and are further emphasized with insets in each graph. *Cloud points were calculated using 4PL parameters which had the best fit for all the samples/assays. *Cloud point calculations: temperature cloud points for the combined AR/ABI1 versus the individual samples were subtracted from the final value to the relative individual values to further emphasize the change using the following formula;* $\Delta Temp = Final_{(AR+ABI1)} - X_{(ABI1\ or\ AR)}$

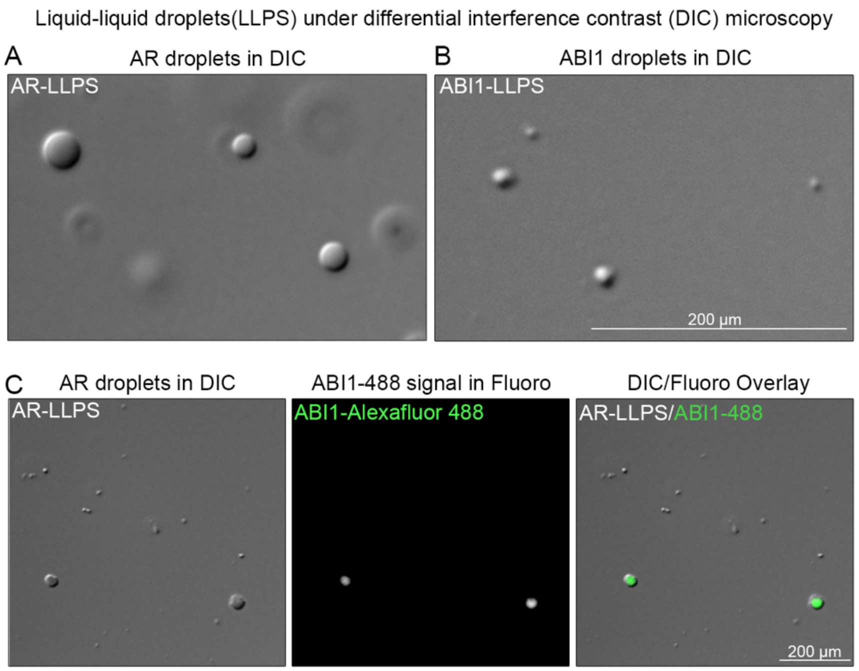


**Supplementary Figure 4. LLPS of AR and ABI1.**

**A-B.** DIC images of *in vitro* LLPS droplets of purified AR (A) and purified ABI1 (B) protein. **C.** Co-phase separation of AR and ABI1: AR LLPS (left panel) and ABI1 (middle panel) shown by ABI1 protein labeled with maleimide Alexa Flour 488 inside LLPS droplets of AR in the same field of view as the overlay shown in (*right panel).* AR LLPS droplets showing propensity for AR to phase separated by itself as shown in (Bouchard et al., 2018)*.* Images were taken using Zeiss Upright AxioImager.Z1 widefield with 100X oil immersion 1.4 NA DIC objective.

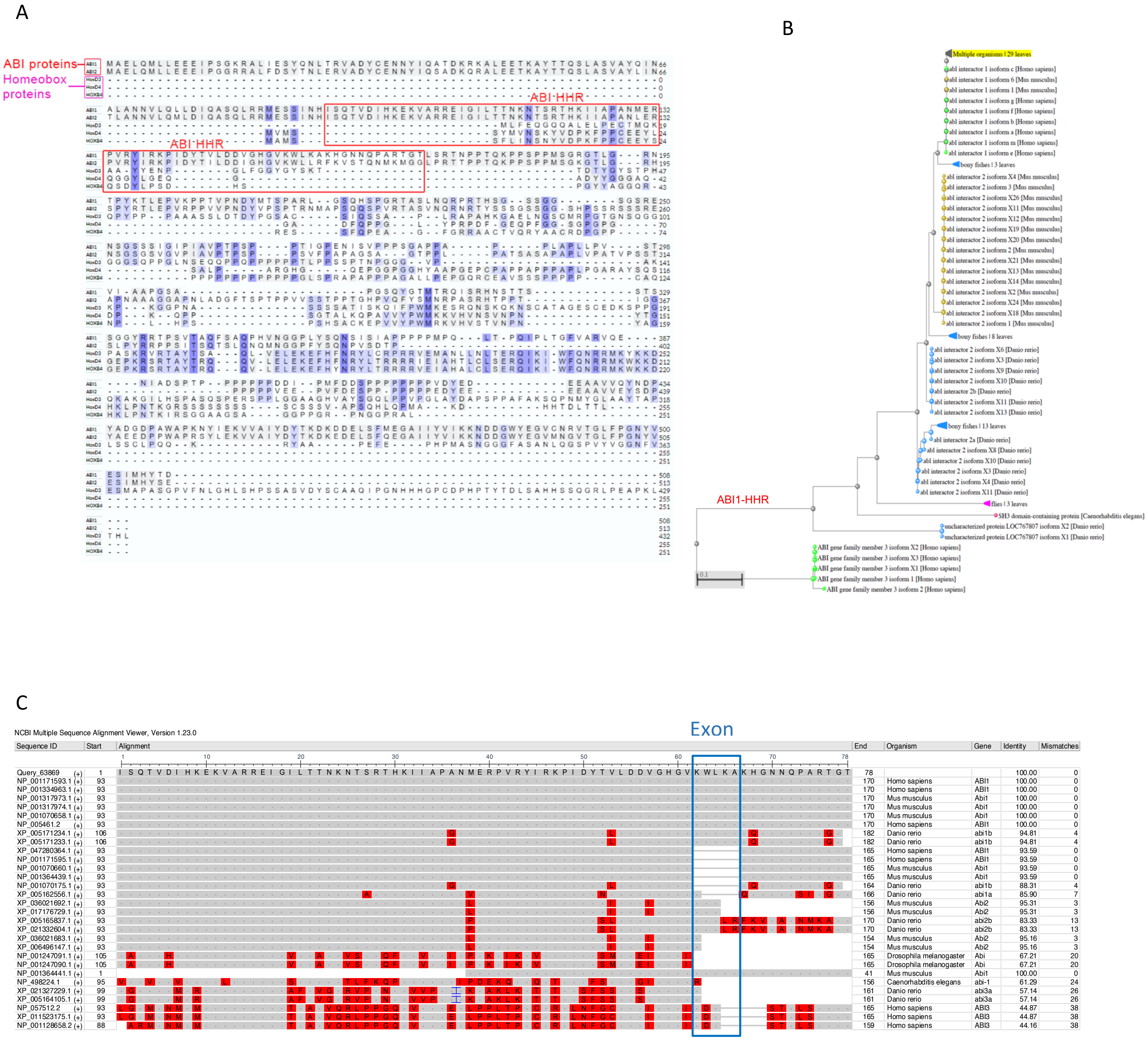


**Supplementary Figure 5. Evolutionary conservation of ABI1-HHR. A. ABI1/2 protein sequence alignment with homeobox proteins.** Alignments were performed using the Unipro alignment feature. Sequences that are aligned include ABI1: ABI1 isoform 1, ABI2 isoform 2; Homeobox proteins: HoxD3, HoxD4, and HoxB4. Classical homeobox proteins are transcription factors which bind highly conserved homeobox DNA sequences. It is evident that the ABI-HHR region does not have high sequence similarity with homeobox proteins which is essential to perform classical homeobox DNA binding functions. ABI1 and ABI2 have high sequence similarity in N-terminal regions of the protein overlapping with the HHR domain. Only human sequences were used for this alignment.

**B-C. Evolutionary conservation of ABI1-HHR domain.** **B**, Evolutionary tree using model organisms as indicated in descriptions next to gene symbols and using ABI1-HHR as query sequence (ABI1-HHR). **C,** Analysis of ABI1-HHR indicates high level of identity across species of the HHR sequences between ABI1 and ABI2, and lower between ABI1 and ABI3. ABI1 is highly conserved from human down to worms (*C. elegans*) and flies (*D. melanogaster*), Blue box indicates alternatively spliced Exon 4 encoded sequences, WLKAK (please note that the alignment program places the C-terminal "K" residue in the N-terminal part of the sequence lacking Exon 4, thus explaining the gap "KWLKA" instead of "WLKAK"). Please note that zebrafish (*Danio rerio*) has proteins containing and lacking Exon 4 sequences. The analysis was performed NCBI BLAST program and display using Multiple Sequence Viewer, version 1.23.00. [7] [8].

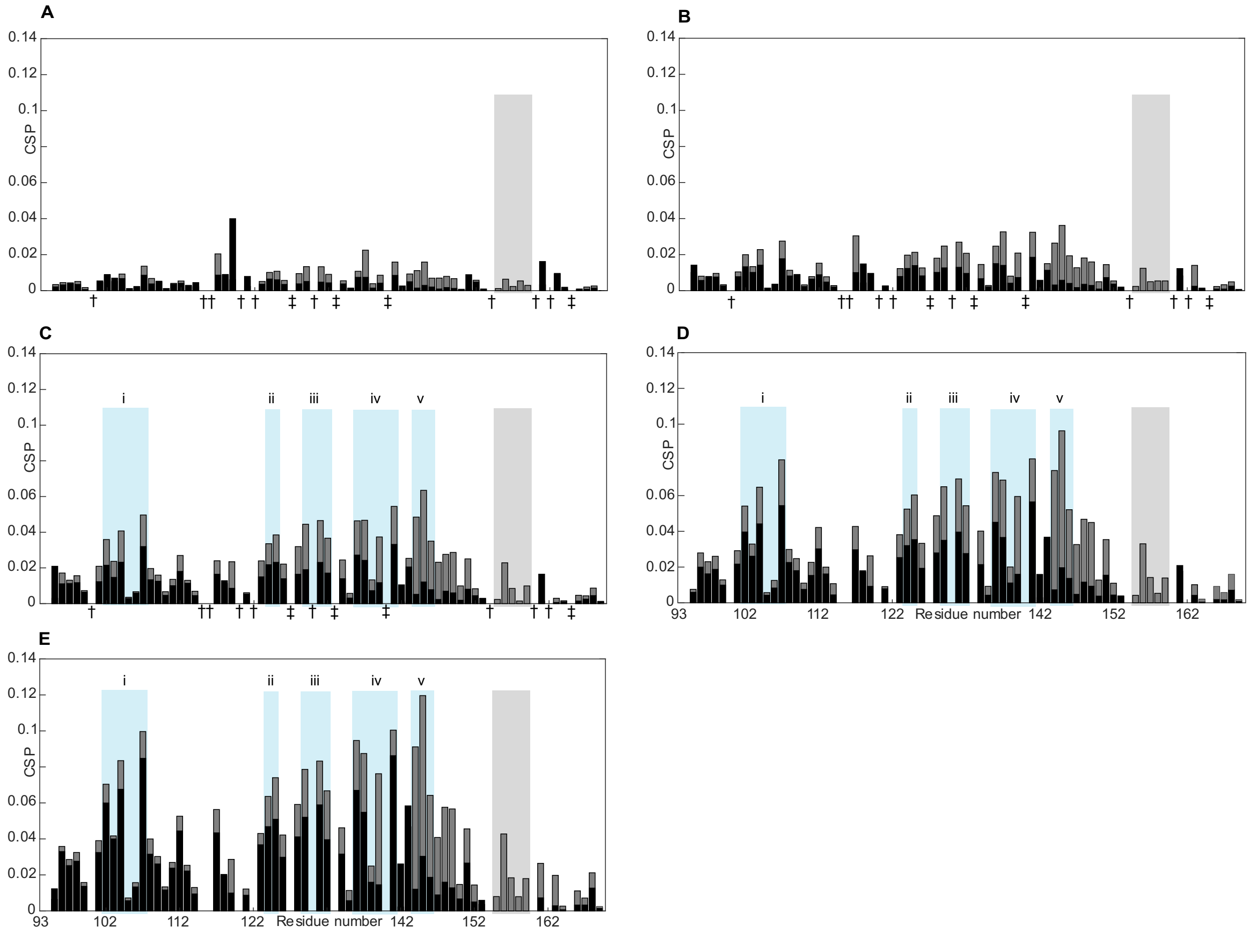


**Supplementary Figure 6. NMR binding analysis for HOXB4 and ABI1-HHR or ABI1-HHR-DelEx4. A-E.** Comparison of Chemical Shift Perturbations (CSPs) of ABI1-HHR (grey) and ABI1-HHR-DelEx4 (black) in presence of HOXB4 at a stoichiometry of **A** 1:0.47, **B** 1:0.94, **C** 1:1.9, **D** 1:3.75, and **E** 1:7.5. Blue rectangles define regions encompassing groups of residues with CSPs higher than one standard deviation from the median CSP in ABI1-HHR. The grey rectangle highlights residues deleted in DelEx4 (155-159). (‡) denote prolines and (†) denote residues whose peaks could not be centered due to overlap in at least one spectrum required for analysis.

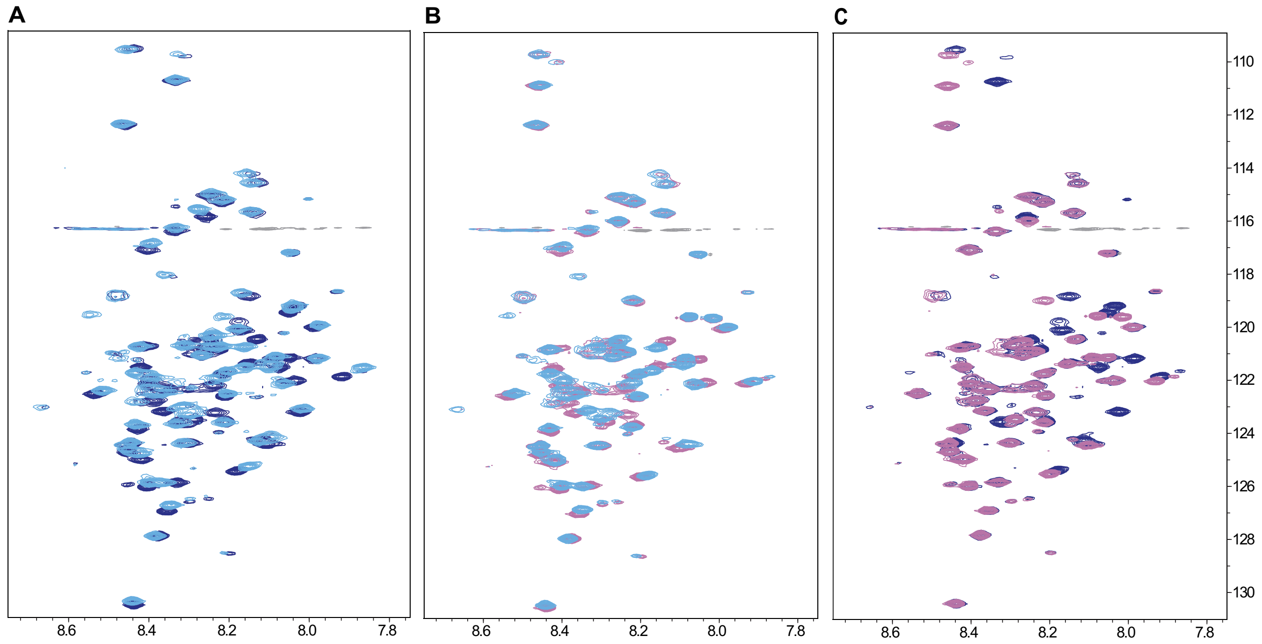


**Supplementary Figure 7. Comparison of ABI1-HHR and ABI1-HHR-DelEx4 NMR spectra with and without HOXB4 and with each other. A-C.** Overlay of $^{1}$H-$^{15}$N HSQC of **A** ABI1-HHR in the absence (dark blue) and presence (light blue) of HOXB4 DNA, **B** ABI1-HHR-DelEx4 in the absence (pink) and presence (light blue) of HOXB4 DNA, and **C** free ABI1-HHR (blue) and free ABI1-HHR-DelEx4. The data shown in **A** and **B** lead to the chemical shift perturbations shown in **Fig. 6E**, whereas the data shown in **C** leads to **Fig. 5L** in the main text.
